# Diet High in Soybean Oil Increases Susceptibility to Colitis in Mice

**DOI:** 10.1101/2021.02.08.430328

**Authors:** Poonamjot Deol, Paul Ruegger, Geoffrey D. Logan, Ali Shawki, Jiang Li, Jonathan D. Mitchell, Jacqueline Yu, Varadh Piamthai, Sarah H. Radi, Kamil Borkowski, John W. Newman, Declan F. McCole, Meera G. Nair, Ansel Hsiao, James Borneman, Frances M. Sladek

## Abstract

The current American diet is high in soybean oil (SO), which consists of unsaturated fatty acids, most notably linoleic acid (LA, C18:2 omega-6). While LA is an essential fatty acid that must be obtained from the diet, high LA consumption has been linked to the development of inflammatory bowel disease (IBD) in humans. Here, we show that a high fat diet (HFD) based on soybean oil increases susceptibility to colitis in wild-type and IL10 knockout mice. It causes immune dysfunction, decreases colon and crypt length and increases intestinal epithelial barrier permeability; these effects were not observed in low LA HFDs. The SO diet also disrupts the balance of isoforms encoded by the IBD susceptibility gene Hepatocyte Nuclear Factor 4α (HNF4α). Both the SO diet and an LA gavage cause gut dysbiosis: the SO diet increases the abundance of an adherent, invasive *Escherichia coli* (AIEC), which can use LA as a carbon source, and the LA gavage decreases the beneficial bacteria *Lactobacillus murinus*. Metabolomic analysis of both host-associated and cultured bacteria shows that SO increases levels of LA and oxylipins while decreasing eicosapentaenoic acid (EPA, C20:5 omega-3) and endocannabinoids. Our results suggest that excess LA, obtained from a diet high in soybean oil, increases susceptibility to colitis by alterations in intestinal HNF4α, gut microbiota and bioactive metabolites.

## INTRODUCTION

Inflammatory bowel disease (IBD) is a multifactorial disorder, the pathogenesis of which can be influenced by host genetics, immune dysfunction, intestinal microbiota, and a variety of environmental factors, including diet (Knight-Sepulveda et al., 2015). Over the last century there has been a shift in the composition of the American diet with soybean oil (SO) being the component that has increased the most (Blasbalg et al., 2011), paralleling the increased incidence of IBD in the U.S. (Molodecky et al., 2012). While the major fatty acid present in soybean oil, linoleic acid (LA, C18:2ω6), has been positively linked to the development of ulcerative colitis (Tjonneland et al., 2009; Wood, 2010), there are conflicting reports on whether soybean oil promotes or protects against IBD (Barros et al., 2010; Moldal et al., 2014; Rashvand et al., 2015; Wiese et al., 2016). Additionally, LA, and its downstream metabolite arachidonic acid (AA, C20:4ω6), are precursors to bioactive lipids such as oxylipins and endocannabinoids (Alvheim et al., 2013; Gabbs et al., 2015). A diet high in LA could thus lead to altered levels of oxylipins and endocannabinoids which have been linked to IBD (Cani et al., 2016; Diab et al., 2019; Zhang et al., 2013).

LA is the endogenous ligand for HNF4α (Yuan et al., 2009), an IBD susceptibility gene (UK IBD Genetics Consortium et al., 2009) and a member of the nuclear receptor superfamily of ligand-dependent transcription factors (Sladek et al., 1990). The human and mouse *HNF4A* genes are highly conserved and contain two promoters (P1 and P2) that drive the expression of P1 isoforms (HNF4α1-6) and P2 isoforms (HNF4α7-12) which have distinct first exons (Ko et al., 2019). While the whole-body knock-out of HNF4α is embryonic lethal (Chen et al., 1994), exon swap mice that express exclusively P1- or P2-derived HNF4α (Briancon and Weiss, 2006) can be used to study the role of HNF4α isoforms in various tissues, including the intestines. Both HNF4α promoters are active in the small intestines and colon, albeit in different parts of the crypt: P1-HNF4α is expressed in the differentiated portion at the top of the colonic crypt and P2-HNF4α in the proliferative compartment in the bottom half of the crypt (Chellappa et al., 2016). While both α1HMZ (expressing only P1-HNF4α) and α7HMZ (expressing only P2-HNF4α) exon-swap mice are healthy under unstressed conditions, we have shown previously that on a low fat, high-fiber diet, α1HMZ males are resistant while α7HMZ males are susceptible to developing dextran sulfate sodium (DSS)-induced colitis (Chellappa et al., 2016).

Intestinal microbiota play an important role in the regulation of gut homeostasis and perturbation in the gut microbiome composition (dysbiosis) can lead to intestinal inflammation and various intestinal pathologies, including IBD (Buttó and Haller, 2016; Ohno, 2015; Tamboli et al., 2004). Notably, increased incidence of the adherent, invasive *Escherichia coli* (AIEC) has been reported in patients with IBD (Darfeuille-Michaud et al., 1998; Palmela et al., 2018), and a Western diet (high in animal fat and sugar) leads to increased intestinal AIEC colonization in genetically susceptible mice (Martinez-Medina et al., 2014). We recently isolated a novel mouse AIEC (*m*AIEC) with 90% DNA sequence homology to the human AIEC and showed that it can cause colitis and exacerbate intestinal inflammation in mice (Shawki et al., 2020).

We have previously shown that a diet high in soybean oil similar to the current American diet can shorten intestinal colonic crypt length (Deol et al., 2015) and that oxylipin metabolites of LA and alpha-linolenic acid (ALA, C18-3ω3) in the liver positively correlate with SO-induced obesity in mice (Deol et al., 2017). While previous studies have demonstrated the role of diet, especially the Western diet, in IBD pathogenesis (Hintze et al., 2018; Marion-Letellier et al., 2016), the effect of the most commonly used cooking oil in the U.S. -- soybean oil -- has not been examined.

In this study, we determined the impact of soybean oil on the development of IBD using a diet that mimics the standard American diet in terms of total fat and LA content. In three different mouse models of colitis -- dextran-sodium sulfate (DSS)-induced, IL10 knockout mice and HNF4α exon swap mice -- we observed increased susceptibility to colitis in adult males fed the SO diet. We examined potential underlying mechanisms including the effect of the diet on HNF4α protein levels as well as the gut microbiome and both host and bacterial metabolomes (see Figure 1 for experimental design). Our results show that while high dietary soybean oil, by itself, does not lead to colitis in wild-type (WT) mice, it does decrease intestinal epithelial barrier function, which could contribute to increased susceptibility to colitis. It also leads to gut dysbiosis as does a daily oral gavage of LA. We show a disruption of the balance between HNF4α isoform levels and an expansion of *m*AIEC in the intestines of SO-fed mice which could also promote susceptibility to colitis. Finally, metabolomic analysis of both host intestinal cells and gut bacteria implicates elevated levels of LA oxylipins as well as reduced levels of endocannabinoids and the omega-3 eicosapentaenoic acid (EPA) in colitis susceptibility. Taken together, our results indicate that a diet enriched in soybean oil, analogous to the current American diet, may be a predisposing environmental factor in the development of IBD and that intestinal barrier dysfunction, HNF4α isoform imbalance, gut bacterial dysbiosis and metabolomic perturbations may contribute to this effect.

**Figure 1.**
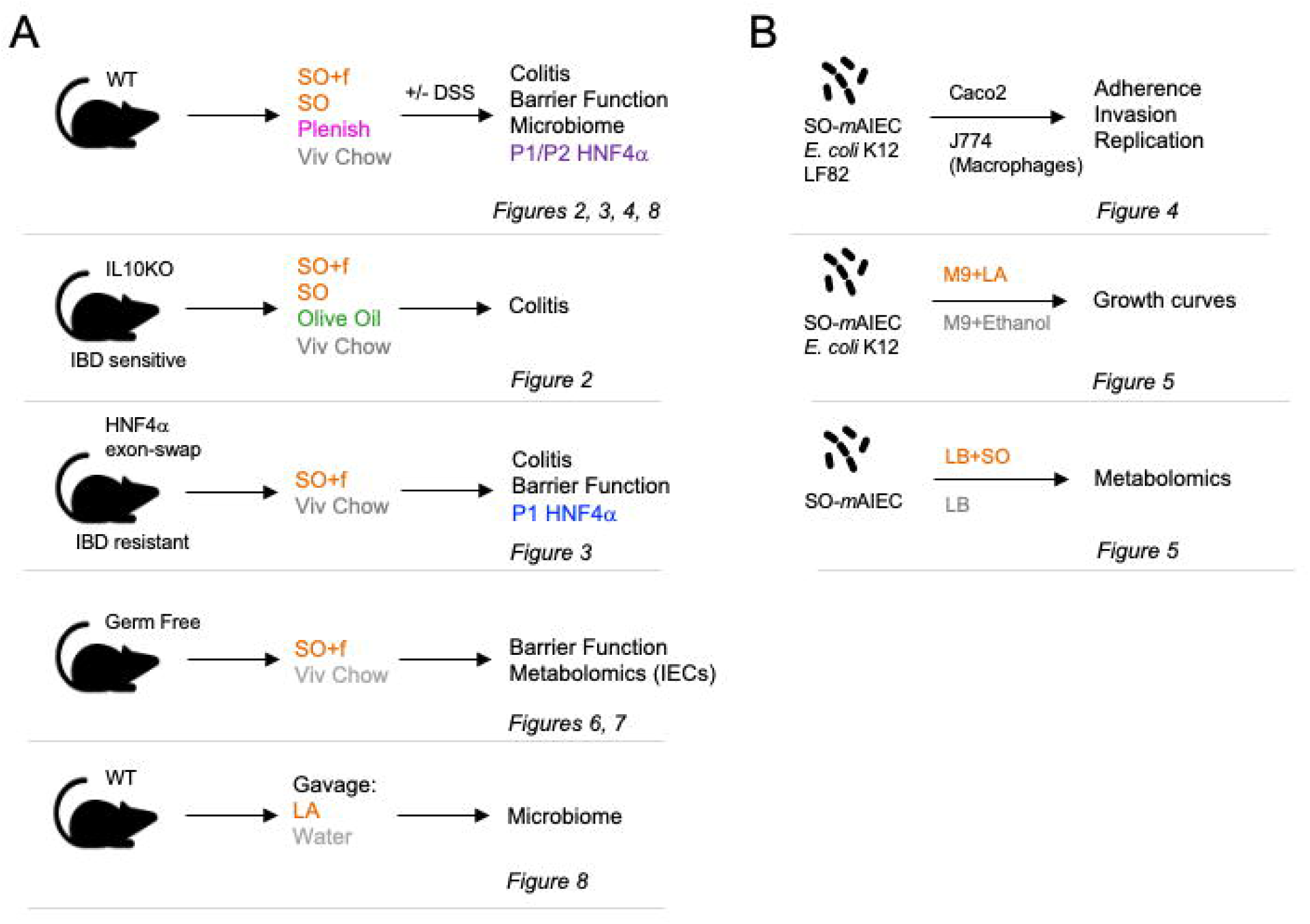
Experimental Design. **(A)** *In vivo* experiments using four different mouse models (C57BL6 adult males) and the indicated treatments with high fat diets (SO+f, soybean oil plus fiber; SO, soybean oil; Plenish, low LA soybean oil and olive oil), DSS and/or gavage with linoleic acid (LA). **(B)** *In vitro* experiments with the indicated bacterial strains and treatment of mammalian cells (CaCO2, J774) or growth condition. The assays used in each model are indicated as well as the figures presenting the results.

## RESULTS

### A diet high in soybean oil increases susceptibility to DSS-induced colitis in WT mice

To determine the effect of soybean oil on susceptibility to colitis, male C57BL6/N mice (WT) were fed either a high fat diet containing soybean oil (35 kcal% fat, 18.6 kcal% LA) or a low-fat vivarium chow (13 kcal% fat, 4.5 kcal% LA). The two diets had similar amounts of fiber: 19% in the SO diet (referred to as SO+f) and 23% in the chow diet (referred to as Viv chow) (Supplementary Table 1). Consistent with our previous studies using a diet high in soybean oil but low in fiber (Deol et al., 2017, 2015), after 15 weeks on the SO+f diet the mice gained significantly more weight than the Viv chow controls (Supplementary Figure 1A). The SO+f mice also had shortened colons (7.9 cm vs 9.5 cm on Viv chow) and crypt lengths (103 µm vs 116 µm on Viv chow) but no morphological differences were apparent in tissue sections from the distal colon (Supplementary Figure 1B). When challenged with DSS after 15 weeks on the diet, we observed a higher percent body weight loss in the SO+f group compared to the Viv chow control starting with day three of the DSS treatment (Figure 2A). Accordingly, the average percent body weight at harvest was significantly lower in the SO+f diet mice (88% vs 98% in Viv chow controls) (Figure 2A).

**Figure 2.**
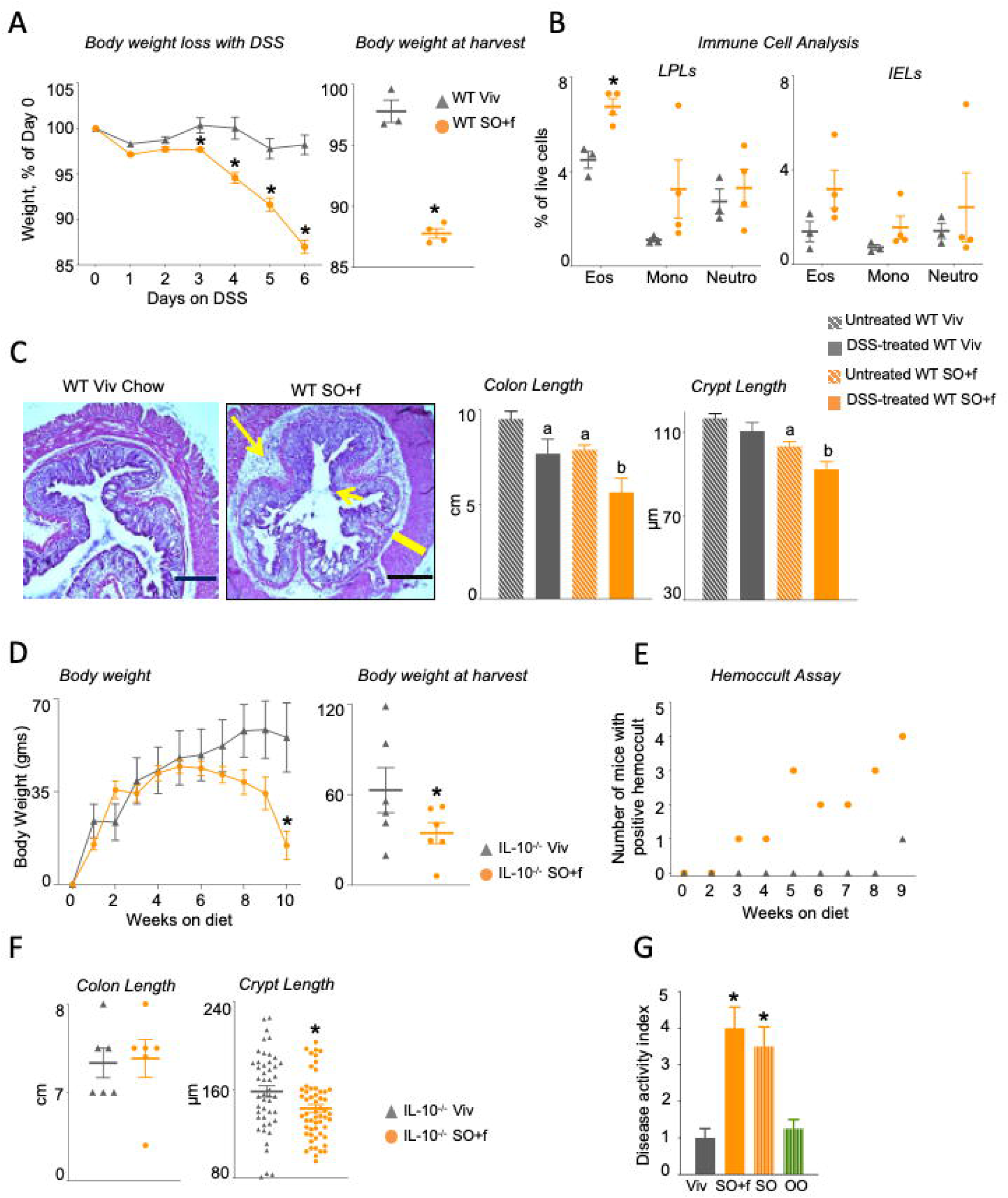
A diet high in soybean oil increases susceptibility to colitis in WT and IL10^-/-^ mice. WT mice on Viv chow or SO+f diet for 15 wks were left untreated or were treated with 2.5% DSS for 6 days: **(A)** % weight loss of DSS-treated mice, body weight at harvest; **(B)** immune cell analysis (Eos, eosinophils; Mono, monocytes; Neutro, neutrophils) in DSS-treated mice, (see Supplementary Figure 1C,D for data from untreated mice); **(C)** colonic histology, colon length and crypt length (at least 10 crypts were measured per mouse) Big arrow, immune infiltrate; small arrow, loss of crypt structure; line, thickening of muscularis in SO+f. Calibration bar = 400 microns. Additional sections are shown in Supplementary Figure 1. IL10^-/-^ mice fed SO+f or Viv diet for 10 wks. **(D)** Average weekly body weights, weight at harvest; **(E)** hemoccult assay; **(F)** Colon length and crypt length. Calibration bar = 400 microns. Colon sections are shown in Supplementary Figure 2. **(G)** Disease activity index (DAI) after 6 weeks on indicated diet. See Supplementary Table 2 for details on DAI scores. * vs Viv, ^a^ vs untreated WT Viv, ^b^ vs untreated WT SO+f. T-test, *P < 0.05* N=5-12 per group.

Since immune dysfunction is an important manifestation of IBD, we characterized immune cell populations in isolated intestinal epithelial lymphocytes (IELs) and lamina propria lymphocytes (LPLs) from both untreated (naive) and DSS-treated Viv chow and SO+f diet mice. Compared to Viv chow-fed mice, a significant increase in the eosinophil population in the intestinal LPLs was observed in SO+f mice after DSS treatment (Figure 2B) but not in untreated mice (Supplementary Figure 1C, D). In contrast, no significant difference was observed between SO+f or Viv chow fed mice for monocytes or neutrophils in the LPLs, nor in any of the immune cell populations in the IELs (Figure 2B). Histological analysis of the distal colon from DSS-treated Viv chow and SO+f diet mice shows that the latter have extensive loss of crypt structure, increased submucosal inflammation and thickened muscularis (Figure 2C and Supplementary Figure 1E). Taken together, these results indicate that DSS treatment causes greater immune dysregulation in mice fed SO+f than in control mice on Viv chow.

DSS treatment caused colon lengths to shorten further in the SO+f group (29% decrease); colon length of the Viv chow mice also decreased compared to the untreated controls, but significantly less so (19% decrease) (Figure 2C, right). Crypt length measurements revealed a similar pattern as colon length, with DSS treatment shortening the crypts in both groups but with only the SO+f mice showing a significant difference (10.5% decrease) (Figure 2C right). As with greater immune dysregulation, decreased colon and crypt lengths are correlated with increased colonic inflammation (Adachi et al., 2006; Lee et al., 2007).

### A diet high in soybean oil accelerates onset of colitis in IL10 deficient mice

To determine whether the soybean oil diet can affect susceptibility to disease in a genetic model of colitis, IL-10 deficient mice (IL-10^-/-^) (B6 strain which develops a mild form of colitis) were fed either the Viv chow or SO+f diet for 10 weeks. IL-10^-/-^ mice on the SO+f diet had accelerated development of disease indices of colitis -- i.e., body weight loss starting at seven weeks and appearance of blood in the stool after three weeks on the diet. In contrast, blood was apparent in the stool of Viv chow-fed animals only after nine weeks and there was no significant drop in body weight even at 10 weeks (Figure 2D, E). Even though both groups of mice had similar weights at the beginning of the dietary treatments, the SO+f diet mice had significantly lower body weights than the Viv chow group at the time of harvest (Figure 2D). This is notable given that the SO+f diet is a high-fat, high-caloric diet. Colon lengths in the two diet groups were similar and there were no obvious differences in gross colonic histology but the SO+f group had significantly shorter crypt lengths compared to the Viv chow control (9.8% decrease) (Figure 2F, right and Supplementary Figure 2B). Compared to WT mice, the crypt length is elongated in IL-10^-/-^ mice on both diets (Figure 2F vs 2C), which is consistent with previous studies (Berg et al., 1996; Lee et al., 2007). No effect was observed on overall small intestinal length by the SO+f diet in IL-10^-/-^ mice (or spleen weight), although the SO+f diet does decrease the liver as a percent body weight (Supplementary Figure 2A).

Finally, to determine whether the added fiber in the SO+f diet (see Supplementary Table 1) plays a role in colitis susceptibility, we fed the IL10 KO mice an isocaloric SO diet (35 kcal% fat) lacking fiber for six weeks and assessed disease activity index (weight loss, ruffled fur, activity, colon weight/length ratio, hemoccult assay, and gross morphological changes) (Supplementary Table 2). The results show nearly identical disease indices between the two SO diets, both of which were significantly greater than the index of the Viv chow fed mice (Figure 2G). Furthermore, to determine whether it is the high caloric content of the SO diet or the soybean oil per se that causes the increased susceptibility, we compared the SO diet to an isocaloric diet consisting of olive oil (OO). After LA, oleic acid (C18:1) is the second most abundant fatty acid in soybean oil – 21% oleic vs 53% linoleic acid – while olive oil consists of 76% oleic acid and 6% linoleic acid, Supplementary Table 1.) The results show that the olive oil diet had a significantly lower disease index than both SO diets and one that was essentially identical to the low-fat Viv chow (Figure 2G).

### A diet high in SO overcomes resistance to DSS-induced colitis in HNF4α exon swap mice

To determine whether soybean oil can impact susceptibility to colitis in a genetic model of IBD resistance, just as it does in the IBD-sensitive IL-10^-/-^ model, we examined the effect of the SO+f diet on DSS-induced colitis in HNF4α exon swap mice which express only the P1 isoform of HNF4α (α1HMZ) (Figure 3A). We have previously shown that α1HMZ male mice exhibit resistance to DSS-induced colitis on Viv chow while mice expressing only the P2 isoforms of HNF4α are highly sensitive to DSS treatment (Chellappa et al., 2016). Interestingly, the α1HMZ male mice fed the SO+f diet did not show significant weight gain compared to the Viv chow group until 19 weeks on the diet versus 10 weeks for the WT mice (Supplementary Figure 3A vs Supplementary Figure 1A). When treated with 2.5% DSS for 6 days, the α1HMZ SO+f diet mice lost significantly more weight (10.3%) than the Viv chow controls (1.7%) (Figure 3B), although the percent weight loss after DSS treatment in SO+f-fed mice was somewhat less in α1HMZ mice than in the WT mice (10.3 vs 13% for WT) (Supplementary Figure 3D).

**Figure 3.**
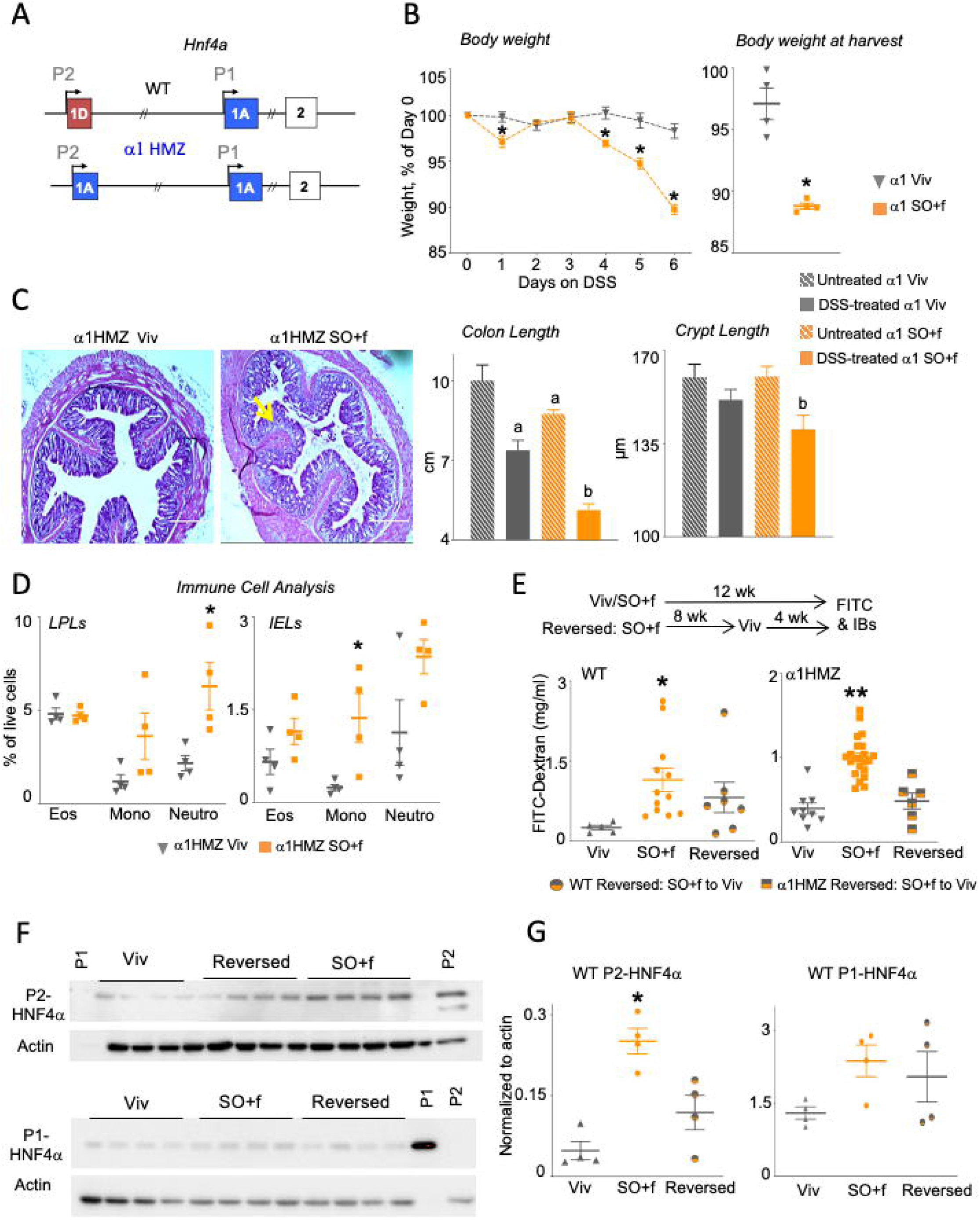
A diet high in soybean oil increases colitis susceptibility in α1HMZ mice and decreases barrier function in WT and α1HMZ mice. (A) Schematic of the mouse *Hnf4a* gene showing the two promoters (P1 and P2) (top), and the exon-swap (1D to 1A) that drives the expression of only P1-HNF4α in α1HMZ (α1) mice (bottom). α1 mice on Viv chow or SO+f diet for 15 wks were treated with 2.5% DSS for 6 days: (B) % weight loss, weight at harvest, * vs α1 Viv *P <* 0.05, T-test N=3-4 per group; (C) colonic histology, colon length and crypt length, ^a^ vs untreated α1 Viv; ^b^ vs untreated α1 SO+f *P <* 0.05 vs other diet, one-way ANOVA, Tukey’s post-hoc. N=3-4 per group. (D) immune cell analysis (Eos, eosinophils; Mono, monocytes; Neutro, neutrophils) in DSS-treated mice, * vs α1 Viv *P <* 0.05, T-test was performed between Viv and SO+f for each cell type, N=3-4 per group. (E) Epithelial barrier permeability in WT and α1 mice fed either Viv or SO+f diets for 12 wks or SO+f for 8 wks, followed by 4 wks of Viv (Reversed) (one outlier removed from WT reversed group). * vs Viv ** vs Viv and Reversed *P* < 0.05, one-way ANOVA, Sidak’s post hoc comparison. N=6-22 per group. (F) HNF4α immunoblots of whole cell extracts (WCE, 30µg) from distal colon of mice fed Viv chow or SO+f diet for 12 weeks or SO+f for 8 weeks followed by Viv chow for 4 weeks (reversed). Each lane contains WCE from a different mouse. P1 control-nuclear extract from HCT116 cells expressing P1-HNF4α; P2 control-nuclear extract from α7HMZ mouse. (See Supplementary Figure 4 for entire blots) (G) Quantification of the P1-and P2-HNF4α immunoblot signals (shown in Fig. 2F) normalized to total protein, as determined by actin staining of the same blot. * *P* < 0.05, one-way ANOVA, Tukey’s post hoc comparison. N=3-4 per group.

As in the WT mice, DSS treatment of α1HMZ mice caused greater loss of crypt structure and more immune cell infiltration in the SO+f diet group compared to the Viv chow control (Figure 3C and Supplementary Figure 3E). At 15 weeks on the diet, the colon length in α1HMZ SO+f fed mice was not significantly different compared to the Viv chow control (8.8 vs 10 cm, *P*= 0.06), nor was there any difference in crypt lengths between the two diets (Figure 3C, right). After DSS treatment, similar to WT mice, colon length in the SO+f α1HMZ mice decreased significantly (from 8.8 to 5.1 cm) and was significantly shorter than the DSS-treated Viv chow group (7.4 cm). DSS treatment also significantly decreased crypt length in SO+f α1HMZ mice compared to the untreated SO+f group (12.5% decrease). However, in α1HMZ mice there was no significant difference in crypt length between Viv chow and SO+f diet either with or without DSS treatment (Figure 3C, right), as there was in WT mice (Figure 2C). Taken together, these results show that while the SO+f diet had less of an impact in α1HMZ compared to WT mice in the absence of DSS treatment, it nonetheless led to increased colitis susceptibility upon DSS treatment even in these “DSS-resistant” mice, confirming again the ability of excess dietary soybean oil to increase susceptibility to colitis.

Immune cell analysis revealed an increase in immune cells after DSS treatment in the α1HMZ mice on the SO+f diet although the profile was different than that observed in WT mice (Figure 3D). While the SO+f diet did not alter the percent of eosinophils in the lamina propria in the α1HMZ mice as it did in WT mice (Figure 2B), there was a significant increase in neutrophils in the LPL population and monocytes in the IEL population with the SO+f diet (Figure 3D). Like the WT mice, there were no significant differences observed in α1HMZ mice in the PBMCs on the two different diets, indicating that the immune dysregulation was localized to the intestine (Supplementary Figure 3C).

### A diet high in soybean oil increases intestinal barrier dysfunction: role for colonic HNF4α isoforms

Having established that the SO+f diet increases susceptibility to colitis in three different mouse models -- DSS WT, IL-10^-/-^ (colitis-sensitive), α1HMZ (colitis-resistant) -- we next investigated the underlying mechanisms by examining intestinal epithelial barrier function using the FITC-Dextran assay. We found that after 12 weeks, the SO+f diet significantly increased epithelial permeability (measured by increased FITC-Dextran in the serum) in both WT and α1HMZ mice (Figure 3E). To determine whether this barrier defect could be reversed by changing the diet, we fed a third group of mice, the SO+f diet for 8 weeks, followed by 4 weeks of Viv chow. We found that while the barrier permeability in WT mice trended lower after the diet reversal, it was not significant. In contrast, in α1HMZ mice, replacing the SO+f diet with Viv chow significantly improved barrier function (Figure 3E).

Immunoblot (IB) analysis of distal colon whole cell extracts showed an increase in the P2-HNF4α isoform in the colons of WT mice fed the SO+f diet (Figure 3F and Supplementary Figure 4A). Changing the diet back to Viv chow did not significantly decrease the P2-HNF4α levels, suggesting that the SO+f diet may cause an irreversible imbalance in the intestinal expression of the HNF4α isoforms (Figure 3F-G, Supplementary Figure 4A). This could explain the lack of significance in terms of rescue of barrier function by Viv chow in WT mice (Figure 3E): we have reported previously that mice expressing only the P2-HNF4α isoform have reduced intestinal epithelial barrier function (Chellappa et al., 2016). In contrast, the level of P1-HNF4α protein in the distal colon of WT (and α1HMZ) mice was not significantly altered by the SO+f diet (Figure 3F-G, Supplementary Figure 4B, C, D). Taken together, these results suggest that a diet high in soybean oil may increase susceptibility to colitis in part by increasing the expression of the P2-HNF4α isoform in the intestines, thereby compromising epithelial barrier function.

### A diet high in soybean oil causes an increase in intestinal *m*AIEC

Another potential mechanism by which a high soybean oil diet may increase susceptibility to colitis is via alteration of the gut microbiome. Examination of the intestinal microbiota revealed that a diet high in soybean oil (with no added fiber, referred to as SO in Supplementary Table 1) causes dysbiosis of the bacteria associated with the intestinal epithelial cells, with a notable increase in the relative abundance of a specific *Escherichia coli* phylotype (Figure 4A). Since a portion of the rRNA ITS region of this *E. coli* phylotype had 100% sequence identity with an AIEC we recently characterized in another genetic mouse model of IBD susceptibility (Shawki et al., 2020), we conducted a phenotypic characterization of the isolate. In Caco2-brush border epithelial cells (Caco-2BBE), the *E. coli* isolate (designated SO-*m*AIEC) demonstrated increased adherence and invasion compared to non-pathogenic *E. coli* K-12. It also had values that were only slightly less than those of the well characterized AIEC LF82, which is associated with IBD in humans (Darfeuille-Michaud et al., 2004). SO-*m*AIEC also showed greater replication in murine macrophages (J774A.1) than both *E. coli* K-12 and AIEC LF82 (Figure 4B): since LF82 is a human isolate, we would not expect it to invade and replicate in mouse macrophages as well as the putative mouse AIEC. On the other hand, since Caco2 is a human-derived cell line, we would expect LF82 to adhere to and invade these cells more effectively than our putative mouse AIEC. These results confirmed that the *E. coli* phylotype and isolate, which increased in abundance in the intestines of mice fed the SO diet, is indeed an AIEC; hence the designation SO-*m*AIEC, the full strain designation being UCR-SoS5.

**Figure 4.**
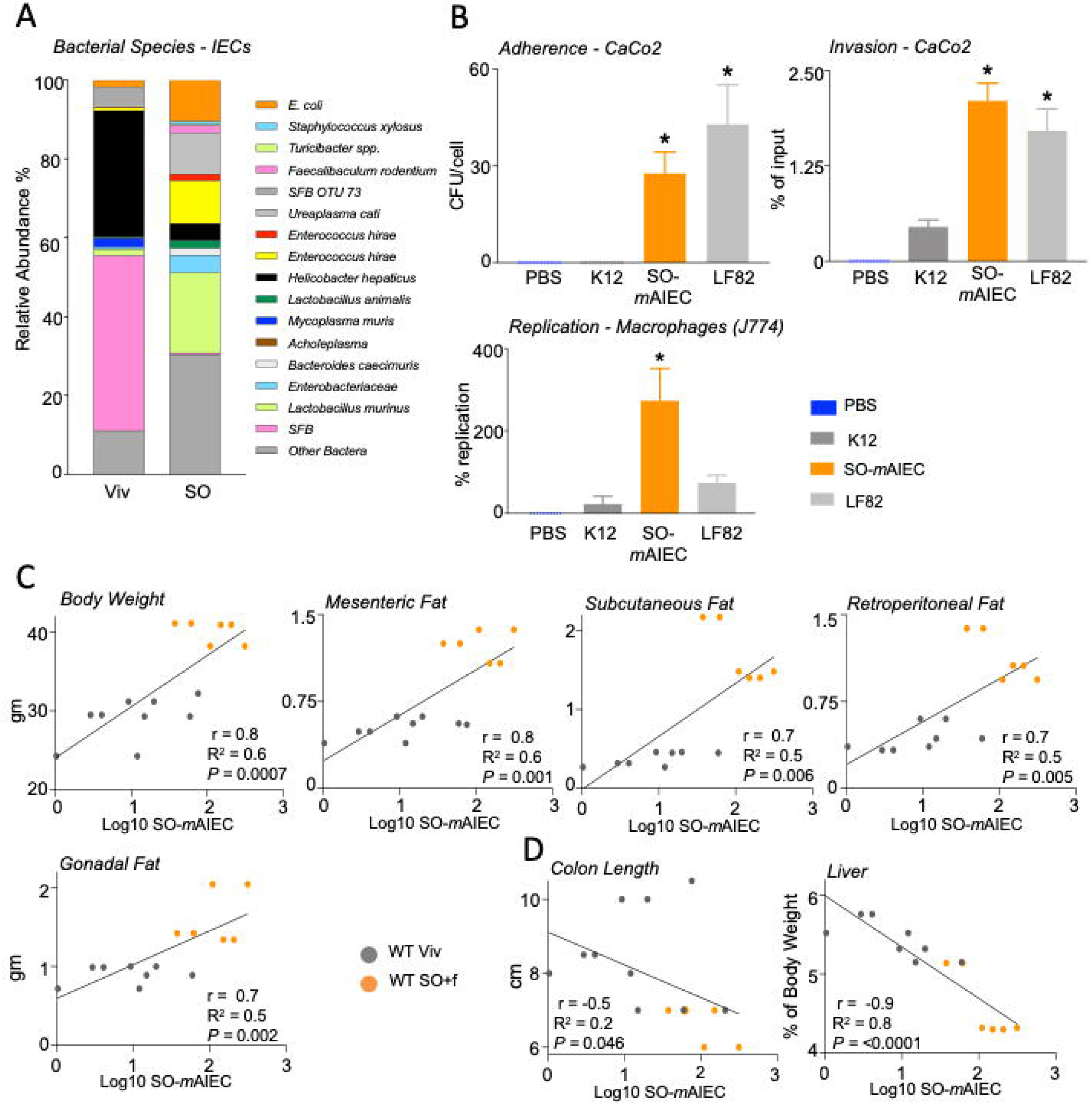
A diet high in soybean oil increases the abundance of SO-*m*AIEC in WT mouse intestines. **(A)** Bacterial species plots of intestinal epithelial cells (IECs) from the small and large intestines (N=4–5) of mice fed Viv chow or an SO diet, with no added fiber. **(B)** Phenotypic characterization of the *E. coli* isolate enriched by SO (SO-*m*AIEC), compared with the human AIEC LF82 and the non-pathogenic *E. coli* K12. Assessments were made for bacterial adherence to Caco-2BBe cells, intracellular invasion of CaCo-2BBe cells, and replication in J774A.1 murine macrophages. * *P* < 0.05, one-way ANOVA, Tukey’s post hoc comparison. N=12 across 4 experiments. Positive **(C)** and Negative **(D)** correlations between indicated mouse metadata and log10 of SO-*m*AIEC relative abundance in the IECs of mice fed Viv or SO+f diets. Pearson correlation coefficient (r) goodness of fit or R^2^ values for linear regression and P value of the correlations (*P)* are indicated on the graphs.

The intestinal abundance of the SO-*m*AIEC directly and significantly correlated with body weight and adipose tissue weight (Figure 4C). Conversely, colon length showed a modest negative correlation with SO-*m*AIEC abundance (Figure 4D), raising the possibility that this bacterium could play a role in the reduction of colon length in mice fed the SO+f diet (Figure 2C). Liver as a percent of body weight also decreased with increasing SO-*m*AIEC levels (Figure 4D), suggesting that the intestinal bacterial dysbiosis induced by the SO diet could potentially have effects outside of the gut.

### SO-*m*AIEC can accumulate, utilize and metabolize linoleic acid

To further investigate the link between a high soybean oil diet and the SO-*m*AIEC, we conducted a targeted quantitative metabolomic analysis of SO-*m*AIEC grown *in vitro* in media (LB) with or without soybean oil; we also examined the metabolomes of both the bacteria and the media (Figure 5A, Supplementary Figure 5 B, C). Since soybean oil is high in polyunsaturated fats (PUFAs: 53% LA and 7% alpha-linolenic acid, ALA, C18:3ω3), we used a platform that measures bioactive metabolites of PUFAs, such as oxylipins and endocannabinoids. Principal components analysis (PCA) of the results shows a notable difference in the profiles obtained for SO-*m*AIEC grown in the absence or presence of soybean oil for five of six samples (Figure 5A). This difference was driven mainly by the levels of the omega-6 LA and the omega-3 ALA and their epoxy-derivatives which coalesced into a single cluster component (Cluster 10), and the omega-3 EPA, which is derived from ALA, and its metabolites (Cluster 12) (Supplementary Figure 5A and Supplementary Table 3). The levels of LA and ALA were increased significantly in SO-*m*AIEC grown in the presence of soybean oil compared to control media (Figure 5B), as were the levels of three oxylipin metabolites of LA (9-KODE, 9,10-EpOME and 12,13-EpOME) (Figure 5C). In contrast to LA, EPA was lower in bacteria grown in the presence of soybean oil (Figure 5B). Similarly, arachidonic acid (AA), a fatty acid precursor for numerous inflammatory mediators derived from LA, did not differ in SO-*m*AIEC grown with or without soybean oil (Supplementary Figure 5B). None of these compounds showed a significant difference in levels in the growth media alone (Supplementary Figure 5C). Taken together, these results indicate that the SO-*m*AIEC isolate can accumulate and metabolize LA from soybean oil.

**Figure 5.**
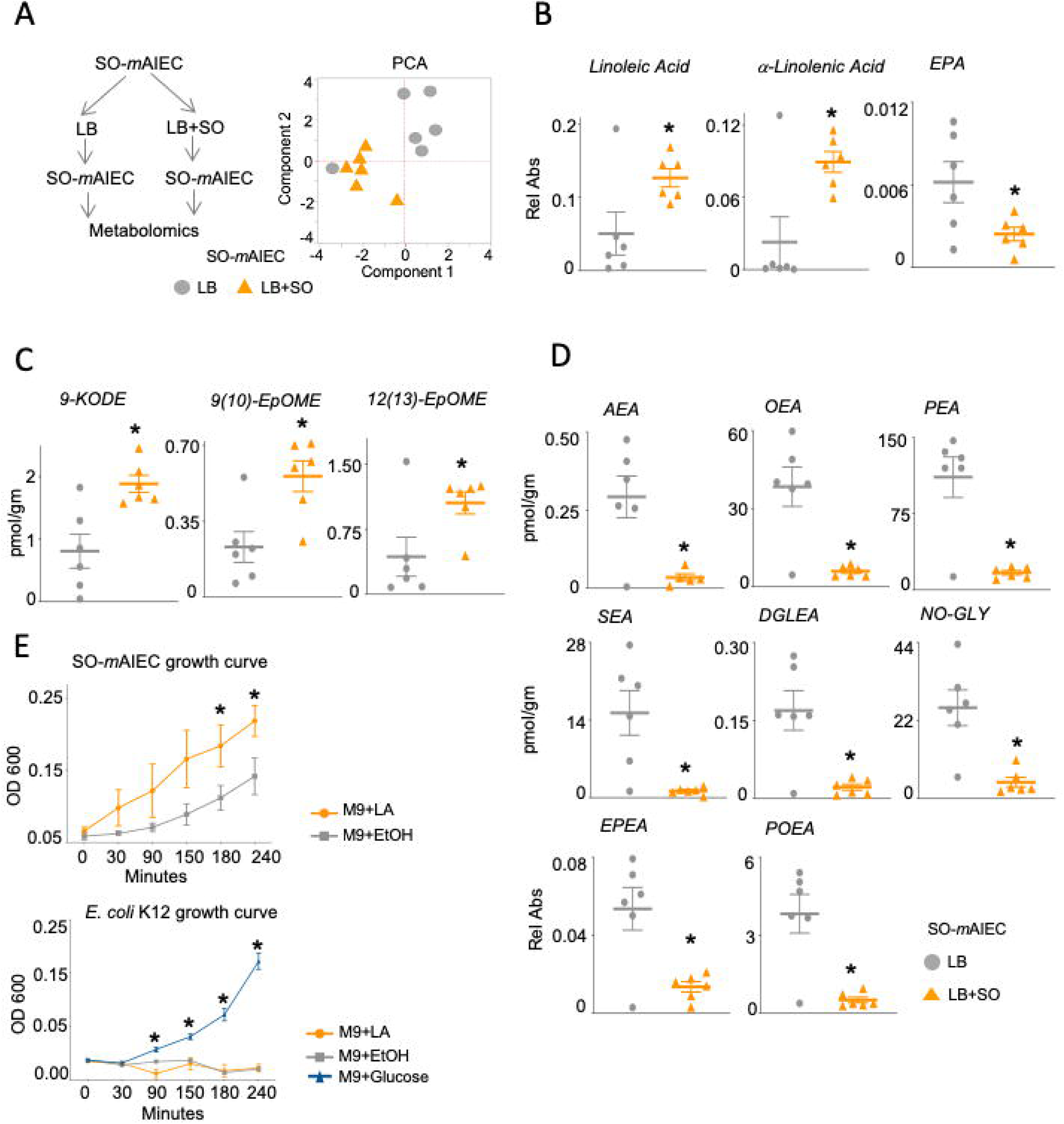
Soybean oil increases oxylipins and decreases endocannabinoids in SO-*m*AIEC cultured *in vitro*. (A) Experimental workflow and Principal Components Analysis (PCA) of oxylipin and endocannabinoid metabolites in SO-*m*AIEC grown with or without SO in the media. Absolute levels of fatty acids (B), oxylipin metabolites (C) and endocannabinoids (D) measured in SO-*m*AIEC grown *in vitro* in the presence or absence of SO in the media. * *P* < 0.05, T-test. N=6 per group. (E) Growth curves for SO-*m*AIEC and *E.coli* K-12 grown in Minimum Essential Medium (M9) with LA, ethanol or glucose as the carbon source. * *P* < 0.05, T-test. N=3 replicates per culture.

Endocannabinoids derived from LA (linoleoyl ethanolamide, LEA), ALA (alpha-linolenoylethanolamide, aLEA) or DHA (docosahexaenoylethanolamine, DHEA), did not show a significant difference in levels in SO-*m*AIEC grown with or without soybean oil (Supplementary Table 4). However, the endocannabinoid anandamide (N*-*arachidonoyl ethanolamide, AEA), which is derived from AA, and closely related N-acyl ethanolamides -- N-oleoyl ethanolamide (OEA), N-palmitoyl ethanolamide (PEA), N-stearoyl ethanolamide (SEA), N-dihomo-γ-linolenoylethanolamide (DGLEA), N-oleyl glycine (NOGLY), eicosapentaenoyl ethanolamide (EPEA) and palmitoleoyl ethanolamide (POEA) are all decreased in SO-*m*AIEC when the growth media is supplemented with soybean oil (Figure 5D). Furthermore, the levels of all the endocannabinoids in the growth media were significantly lower than in the bacteria themselves (Supplementary Figure 5C versus Figure 5D), suggesting that while the SO-*m*AIEC can produce these bioactive compounds, their production is blocked by the presence of soybean oil.

To determine whether the SO-*m*AIEC can utilize LA as an energy source, we grew SO-*m*AIEC in a minimal medium (M9) supplemented with either ethanol or LA as the sole source of carbon. The SO-*m*AIEC grew significantly faster in the presence of LA, indicating that it can use LA as a carbon source; in contrast, the non-pathogenic *E.coli* K12 cannot use LA as a carbon source (Figure 5E). Addition of glucose to the LA culture confirmed the viability of the initial inoculate and showed that LA may be lethal to *E.coli* K12 (Supplementary Figure 5D), consistent with published results indicating that LA is bacteriostatic to a number of microorganisms, including the probiotic *Lactobacillus* species (Nieman, 1954; Zheng et al., 2005).

### Intestinal epithelial barrier function in germ-free mice is affected by the soybean oil diet

Our results thus far suggest that dietary soybean oil may induce susceptibility to colitis by providing an environment conducive for the outgrowth of the pathobiont SO-*m*AIEC, which in turn increases the levels of LA oxylipins and decreases the levels of endocannabinoids in the gut. To determine whether the host can also contribute to alterations in the gut metabolome, we employed a germ-free (GF) mouse model. GF animals are known to have a reduced mucus layer due to the absence of bacteria (Schroeder, 2019) and hence are extremely sensitive to epithelial injury during DSS treatment (Hernández-Chirlaque et al., 2016). Therefore, we examined intestinal epithelial barrier function in both GF and conventionally raised (Conv) mice using the FITC Dextran assay as an indicator of IBD susceptibility (Martini et al., 2017).

Consistent with the results in Figure 3E, the SO+f diet significantly increased barrier permeability in Conv mice fed SO+f for 12 weeks versus the Viv chow control (Figure 6A). While the difference in barrier permeability between GF mice on Viv chow versus SO+f diet was not significant, FITC levels in the serum of GF SO+f mice were significantly (*P* < 0.001) greater than in the Conv SO+f mice (Figure 6A). Although elevated, barrier permeability in GF mice fed Viv chow compared to Conv mice was not significant, *P* = 0.06. Taken together, these results suggest that while the presence of gut bacteria in Conv mice is important for decreased barrier function induced by the SO+f diet, it is also possible that some component, or host-derived metabolite, of the SO+f diet may exacerbate barrier dysfunction in GF mice.

**Figure 6.**
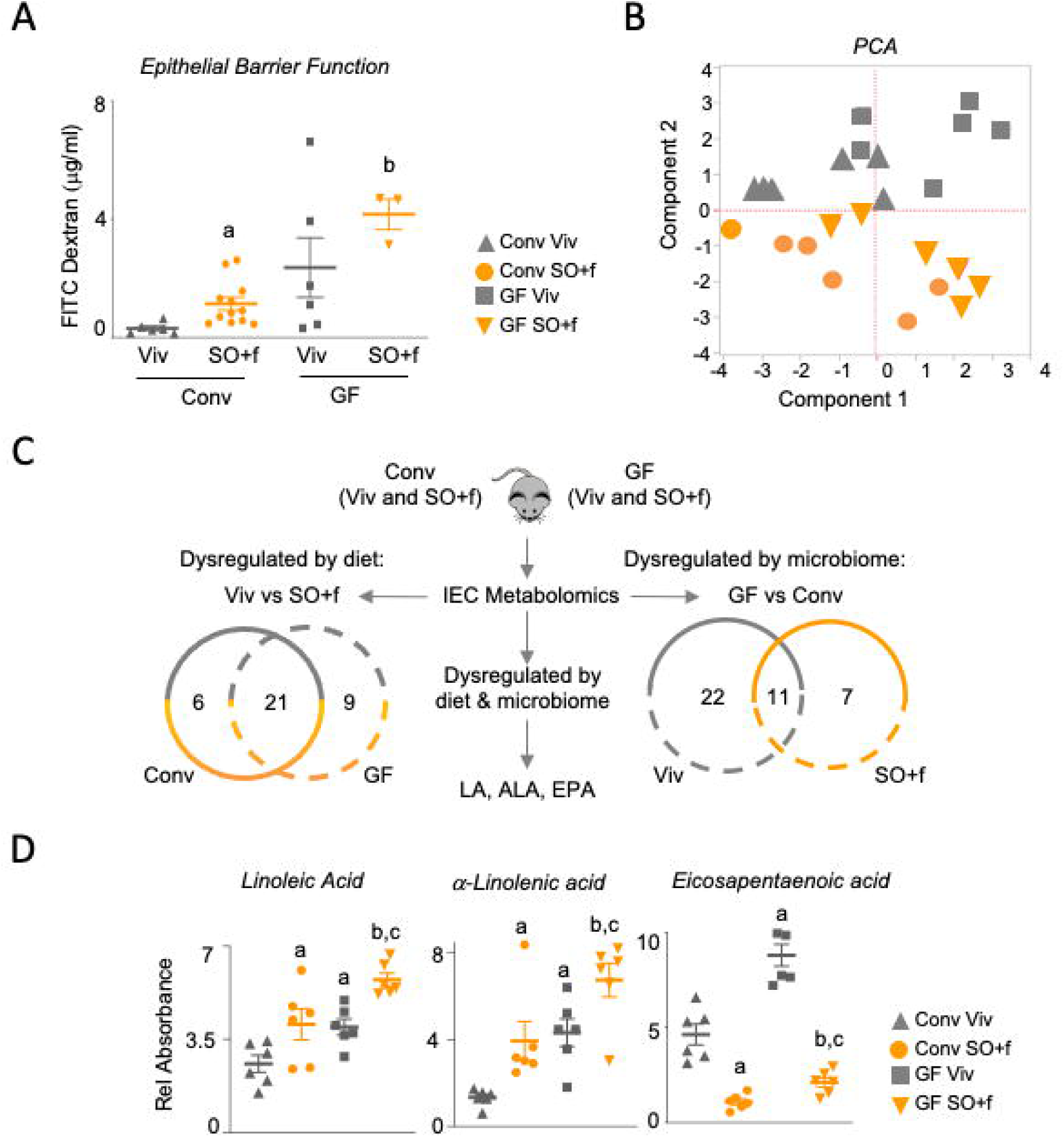
A diet high in soybean oil decreases barrier function and alters the metabolome in the intestines of conventionally raised and germ-free mice. (A) Epithelial barrier permeability in Conv and GF mice fed either Viv chow or SO+f diet for 12 wks. N=3-13 per group. ^a^ *P* < 0.05 vs Conv Viv, ^b^ *P* < 0.001 GF SO+f vs Conv SO+f, *P*=0.06 Conv Viv vs GF Viv, T-test. N=3-13 per group. (B) PCA of oxylipin and endocannabinoid metabolites in IECs isolated from Conv and GF mice fed either Viv chow or SO+f diet for 8 wks. N= 6 per group. (C) Schematic and Venn analysis of metabolomics data in B. (D) Absolute levels of fatty acids measured in IECs from Conv and GF mice fed either Viv chow or SO+f diets for 8 wks. ^a^ vs Conv Viv, ^b^ vs Conv SO+f, ^c^ vs GF Viv *P* ≤ 0.05, T-test. N=6 per group.

To determine which metabolites might be involved in compromising the barrier function we applied the same metabolomic analysis as in Figure 5 to the IECs isolated from Conv and GF mice fed either Viv chow or the SO+f diet. We chose a shorter time on the diet (eight weeks) to identify compounds that might cause barrier dysfunction without the confounding factor of excessive adiposity seen at later time points. At eight weeks on the diet, Conv mice on the SO+f diet were just beginning to increase in body weight and serum FITC; in contrast, there was no difference in body weight between the GF Viv chow and SO+f mice (Supplementary Figure 6A vs Figure 6A, Supplementary Figure 6B). This is consistent with previous reports showing that GF mice are resistant to HFD-induced obesity (Bäckhed et al., 2007; Turnbaugh et al., 2008). PCA analysis of the metabolites shows that the two dietary groups (SO+f and Viv chow) are separated from each other regardless of the microbiome status (Conv or GF) of the mice, reinforcing the notion of host-dependent effects (Figure 6B). Variables with the largest contribution to this separation were EPA and docosahexaenoic acid (DHA) and their lipoxygenase and soluble epoxide hydrolase derivatives (Cluster 1) (Supplementary Figure 6C and Supplementary Table 3).

To better decipher the contributions of diet and the microbiota to the gut metabolome *in vivo*, we conducted a comparative analysis of the metabolites that were significantly dysregulated in IECs harvested from the various conditions: Viv chow vs SO+f diet and Conv vs GF mice (Figure 6C). Analysis of metabolites dysregulated by diet identified a total of 27 oxylipins that were significantly dysregulated in the IECs isolated from Conv mice fed Viv chow versus the SO+f diet. In the GF comparison, 30 oxylipins differed between the Viv chow and SO+f diet fed mice (Figure 6C). Of these, only four are also significantly altered in SO-*m*AIEC in the SO media while 21 of these compounds are commonly dysregulated in both Conv and GF comparisons (and in the same direction), suggesting that diet and/or host cells rather than bacteria play a role in their accumulation/metabolism (Figure 6C, Supplementary Figure 7).

Analysis of metabolites whose levels are significantly altered by the microbiome (Conv to GF comparison for both Viv chow and SO+f diets) identified 33 metabolites that differ in the Viv chow comparison and 18 in the SO+f comparison (Figure 6C and Supplementary Figure 8). Of these, 11 are common to both GF and Conv mice, suggesting that the presence or absence of gut microbiota is the determining factor for these metabolites rather than diet. LA, ALA and EPA were the only three compounds dysregulated in both the Viv chow versus SO+f (21 common metabolites) and the GF versus Conv comparisons (11 common metabolites) (Figure 6C, D). In both GF and Conv mice, LA and ALA were increased by SO+f while EPA was decreased, reminiscent of the *in vitro* results with SO-*m*AIEC in the SO-supplemented growth media (Figure 5B). Intriguingly, all three compounds were increased in GF Viv chow mice compared to Conv mice suggesting that in addition to a diet, bacteria also impact the level of these fatty acids.

In contrast, host cells rather than bacteria appear to play a primary role in the accumulation/metabolism of the LA-derived oxylipin 12,13-DiHOME and the ALA-derived oxylipin 12,13-DiHODE as both Conv and GF mice fed the SO+f diet show decreased levels of these metabolites in the IECs (Figure 7A). In Conv mice, two oxidized LA metabolites, 13-HODE and 13-KODE, and the ALA-derived 13-HOTE are increased in the SO+f fed group (Figure 7A). 13-HODE and 13-KODE are also increased in GF mice fed Viv chow vs the Conv Viv group but none of these are changed in the GF Viv vs SO+f comparison suggesting that bacteria alone could be a factor in their accumulation, but that diet is a factor only in the presence of bacteria.

**Figure 7.**
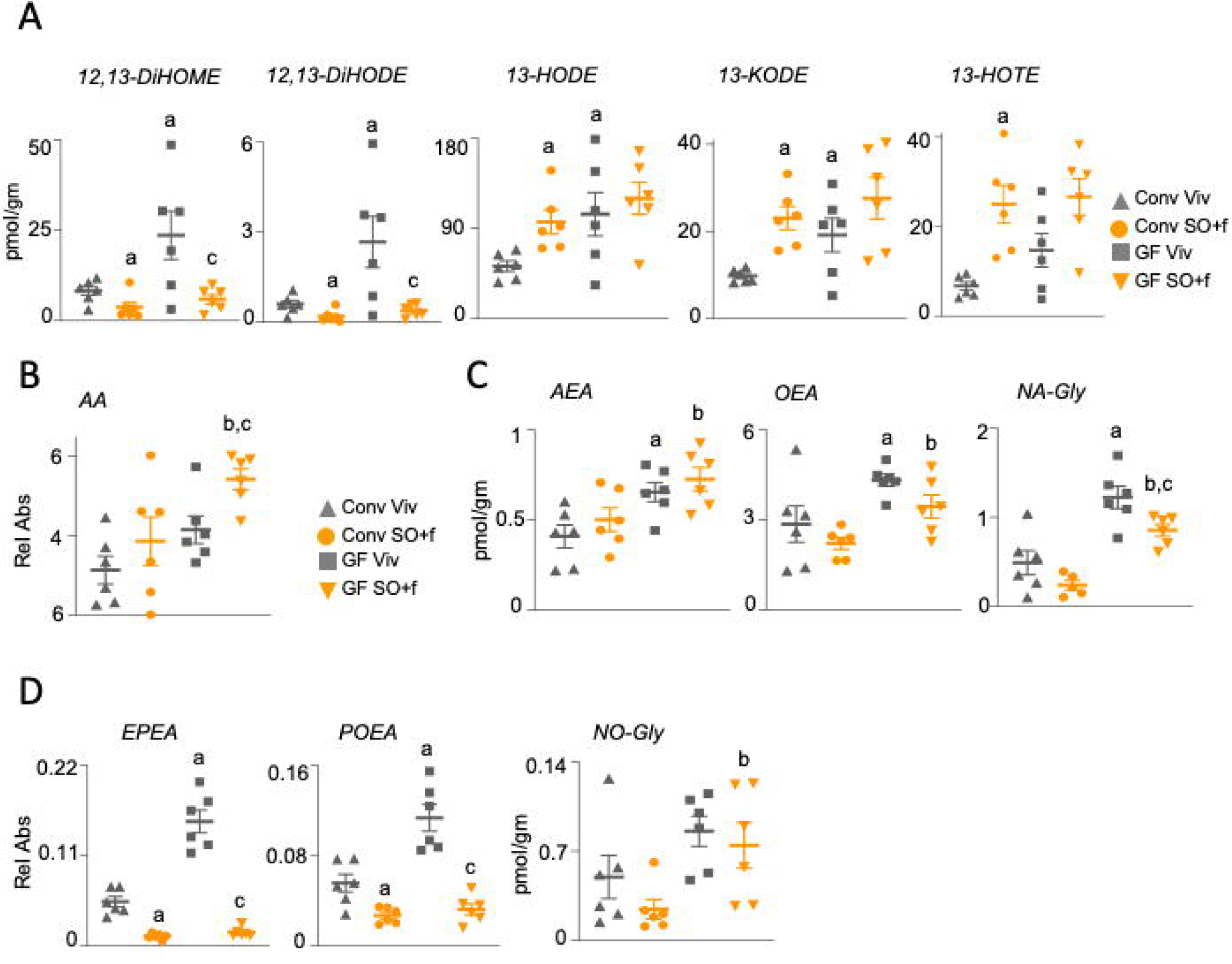
A diet high in soybean oil alters the levels of oxylipin metabolites and endocannabinoids in conventionally raised and germ-free mice. Absolute levels of oxylipins. (A), arachidonic acid (AA) (B) and endocannabinoids (C, D) measured in IECs from Conv and GF mice fed either Viv chow or SO+f diet for 8 wks. ^a^ vs Conv Viv, ^b^ vs Conv SO+f, ^c^ vs GF Viv, *P* < 0.05; *P* =0.07 for AA Conv Viv vs GF Viv, T-test. N=6 per group.

AA levels were significantly higher in GF mice fed the SO+f diet compared to their Viv chow counterparts, as well as the Conv mice on the SO+f diet (Figure 7B) indicating that AA levels can be affected by both diet/host and microbiome status. AA-derived endocannabinoids were similarly dysregulated by both gut microbiota and diet (Figure 7C, D). Of these, AEA, OEA and NA-Gly are decreased by the presence of microbes in the Conv mice, regardless of diet; the SO+f diet also decreased NA-Gly in GF mice (Figure 7C). EPEA and POEA are decreased by the SO+f diet in both GF and Conv mice, and by the presence of microbes in Conv mice (Figure 7D). In contrast, NO-Gly is increased by soybean oil in the absence of microbes, indicating significant dysregulation by both diet and lack of gut microbes (Figure 7D).

To determine whether the enhanced susceptibility to DSS-induced colitis is specific to the SO diet and LA, we examined the response of WT mice fed a genetically modified SO diet low in LA (Plenish, 2.6% LA versus 19% LA in SO, Supplementary Table 1). The results show that Plenish-fed mice lost less weight and had fewer morphological changes including less immune infiltrate and crypt cell damage in the colon than the SO-fed mice (Figure 8A), suggesting less of an impact on colitis susceptibility than the SO diet. These results are consistent with the reduced disease activity of the IL10 KO mice fed an olive oil diet (Figure 2G), which has a fatty acid composition similar to that of Plenish (Supplementary Table 1). Finally, we gavaged mice with similar amounts of LA as would be ingested with the SO+f diet for three days. Beta diversity analysis of fecal samples showed that the bacterial communities resulting from the LA gavage were distinct from a water gavage, with a notable decrease observed in the relative abundance of *Lactobacillus murinus* (Figure 8B).

**Figure 8.**
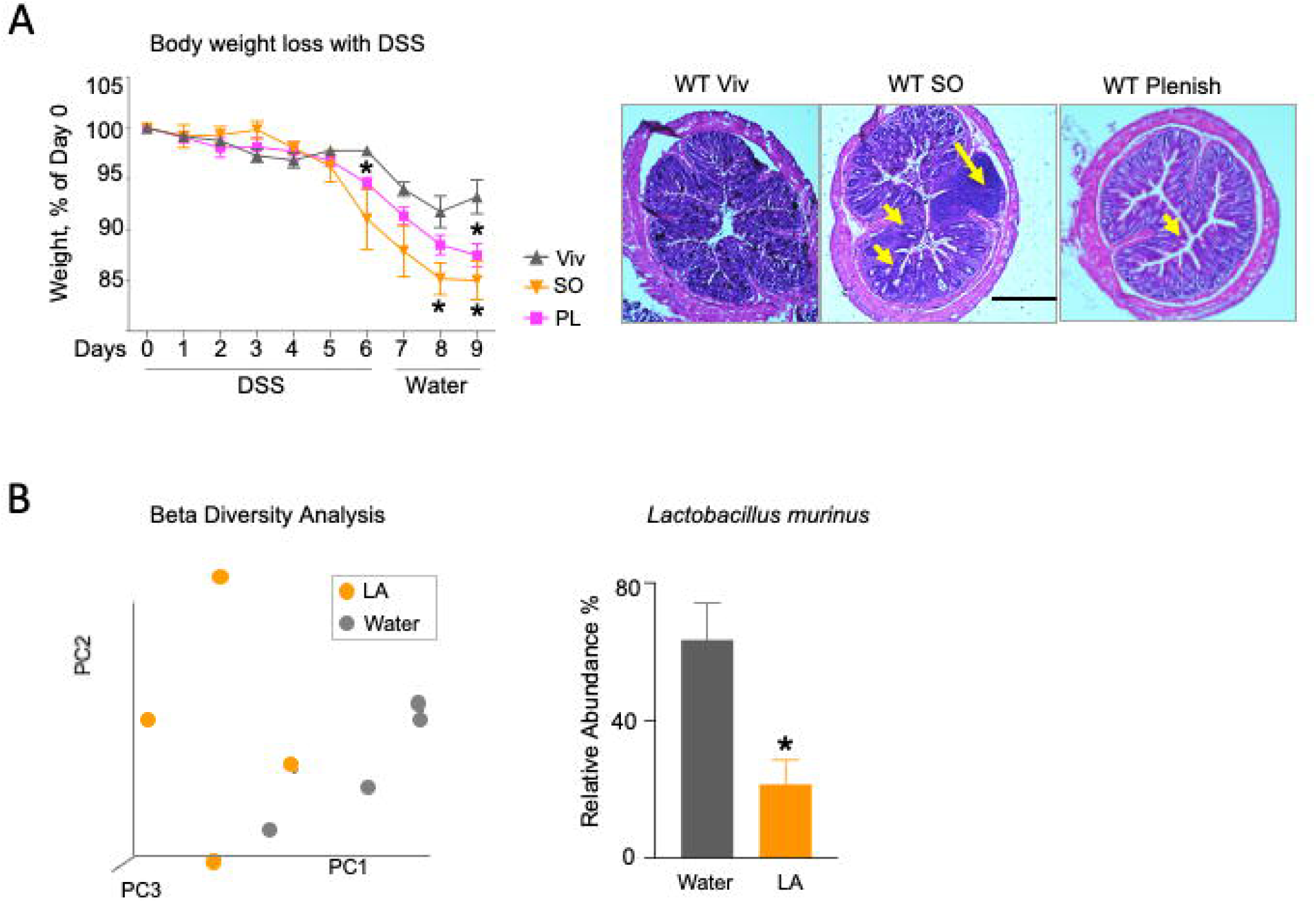
Impact of LA on gut microbiome. **(A)** WT mice on Viv chow, SO or PL diets for 12 wks were treated with 2.5% DSS for 6 days, followed by 3 days recovery on water: *Left panel,* **%** weight loss of DSS-treated mice. *Right panel,* colonic histology. Big arrow, immune infiltrate; small arrow, loss of crypt structure; Calibration bar = 400 microns. Additional sections are shown in Supplementary Figure 9. **(B)** L*eft panel,* bacterial community analysis. Beta diversity distance values were generated from bacterial rRNA ITS sequences and displayed using PCA. *P* = 0.045; PERMANOVA test. N=4 per group. *Right panel,* mean relative abundance of *Lactobacillus murinus* in water or LA gavaged mice. * *P* < 0.05, T-test. N=4 per group.

## DISCUSSION

While the Western diet, high in saturated fat and sugar, has been implicated in the pathophysiology of IBD (Knight-Sepulveda et al., 2015), the role of diets high in unsaturated fats, such as linoleic acid (LA), is less well studied. Soybean oil, ∼55% LA, is the most highly consumed edible oil in the U.S.; its increase in consumption parallels the rise in IBD incidence in humans (Molodecky et al., 2012; USDA Economic Research Service, 2020). In this study we found that a diet high in soybean oil, comparable to the American diet in terms of amount of total fat (35 kcal%), can increase susceptibility to colitis in mice via multiple mechanisms.

Using three different mouse models of colitis -- DSS-induced, IL-10^-/-^ and HNF4α transgenic mice -- we demonstrate that ingestion of amounts of dietary soybean oil comparable to that in the American diet increases intestinal inflammation and modulates the gut microbiome and metabolome, as well as protein levels of the IBD susceptibility gene *Hnf4a* (Figure 9). Specifically, we found that in male mice the soybean oil diet: i) increases intestinal epithelial barrier permeability; ii) increases the abundance of SO-*m*AIEC, a pathobiont that has been shown to cause colitis in mice (Shawki et al., 2020); iii) increases the level of P2-HNF4α in the colon, which is associated with increased sensitivity to DSS (Chellappa et al., 2016); iv) increases putative pro-inflammatory PUFA metabolites, oxylipins; and v) decreases levels of anti-inflammatory bioactive lipids, including endocannabinoids and the omega-3 EPA. Our results indicate that fiber is not a major factor in these effects: unlike most high fat diet studies, the fiber in our soybean oil diet (SO+f) was comparable to that in the low-fat, high-fiber control diet (Viv chow). Furthermore, similar effects of the soybean oil diet were observed in the absence of fiber (Figure 2G).

**Figure 9.**
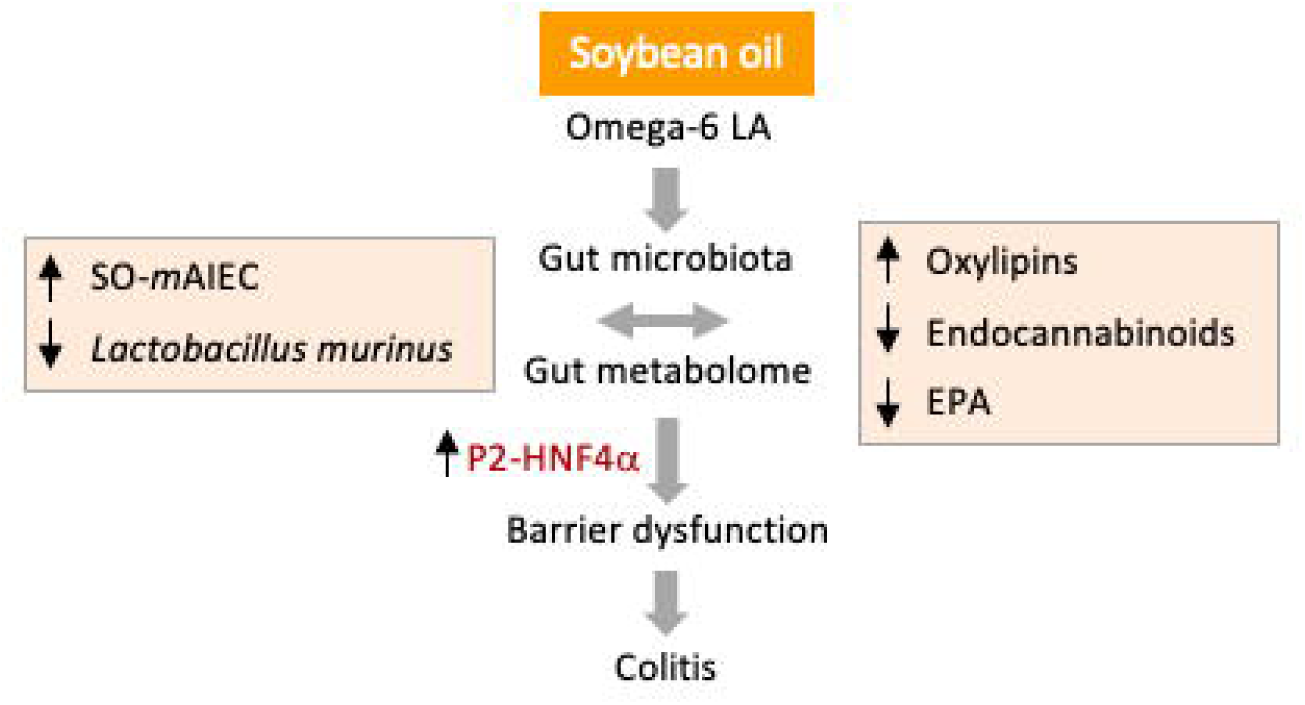
Proposed model by which a diet high in soybean oil increases susceptibility to colitis. See Discussion for details.

Reduction in the diversity of gut bacterial species and an increase in host-associated microbes has been described not only in mouse models of colitis but also in patients suffering from the disease (Gkouskou et al., 2014; Kaur et al., 2011). Beneficial bacteria are decreased in IBD while microbial pathobionts such as AIEC are increased (Alam et al., 2020; Darfeuille-Michaud et al., 2004). The first AIEC strain (LF-82) associated with human IBD was isolated from a patient with Crohn’s Disease (Darfeuille-Michaud et al., 2004). We recently identified a novel mouse AIEC (*m*AIEC) strain with adherence, invasive and replication properties comparable to LF-82 (Shawki et al., 2020). In this study, we report that a diet high in soybean oil increases the population of a related *m*AIEC in the mouse intestinal tract; we refer to it as SO-*m*AIEC. The outgrowth of SO-*m*AIEC could be due to the fact that it can use LA, a major component of soybean oil, as an energy source (Figure 5).

Not unexpectedly, levels of LA as well as ALA (also found in soybean oil although at lower levels than LA) increased in the intestinal epithelial cells (IECs) of both germ-free (GF) and conventional (Conv) mice fed the SO+f diet (Figure 6). Less anticipated was the increased level of these PUFAs in GF mice compared to Conv mice regardless of diet (Figure 6). The *in vitro* results from the SO-*m*AIEC, which had elevated levels of LA, ALA and their oxylipin metabolites in the presence of soybean oil, offer an explanation: gut microbiota can evidently accumulate and metabolize LA and ALA so when they are absent, as in the GF mice, the levels of these PUFAs are elevated in the host.

Neither the olive oil nor the low LA soybean oil (Plenish) diet resulted in the negative effects on colonic health observed with the isocaloric soybean oil diet (Figures 2G, 8A). This suggests that the increased susceptibility to colitis caused by the high soybean oil diet is not due solely to the increase in dietary fat but could be a function of the high LA content and/or LA-derived oxylipins. In fact, we reported previously that a HFD based on Plenish does not lead to elevated levels of oxylipins in the liver or the plasma (Deol et al., 2017). Finally, results from an LA gavage experiment (Figure 8B) support the notion that high levels of LA in soybean oil may play an important role in the increased susceptibility to colitis by decreasing the abundance of beneficial bacteria such as *Lactobacillus murinus* which are protective against colitis and inflammation (Pan et al., 2018; Tang et al., 2015). The toxic effect of LA on various *Lactobacili* has been reported earlier (Nieman, 1954; Zheng et al., 2005). Interestingly, it has been shown that *Lactobacillus reuteri* can develop resistance to LA-induced toxicity in the small intestines (Di Rienzi et al., 2018).

### Role of HNF4**α** isoforms in soybean oil-induced susceptibility to colitis

We have previously shown that increased intestinal expression of P2-HNF4α leads to decreased barrier function and increased DSS-induced colitis (Chellappa et al., 2016). Here, we report that the SO+f diet increased epithelial permeability as well as P2-HNF4α protein (Figure 3), suggesting that an imbalance of the HNF4α isoforms in the gut may play a role in SO-induced colitis susceptibility (Figure 9). While the cause of the increase in expression of P2-HNF4α remains to be determined, bacterial dysbiosis caused by large amounts of soybean oil in the diet could be involved as HNF4α expression has been shown to be modulated by the microbiome (Davison et al., 2017; Lickwar et al., 2022), although the different isoforms were not examined in that study. Another possibility is that LA alters the balance of the HNF4α isoforms. We previously identified LA as the endogenous ligand for HNF4α and showed that it can decrease its transcriptional activity and protein stability (Yuan et al., 2009). However, since both P1- and P2-HNF4α contain identical ligand binding domains, any potential impact of LA on the HNF4α isoform balance would likely be a complex one.

### Role of increased oxylipin levels in SO-induced susceptibility to colitis

LA and ALA oxylipins are bioactive, pro-inflammatory molecules that have been recently linked to IBD (Diab et al., 2019; Marton et al., 2019). Increased levels of these compounds in the intestines of SO-fed mice is reminiscent of our previous findings showing that LA and ALA oxylipin levels in the liver correlate with obesity, another condition linked to chronic inflammation (Deol et al., 2017; Ellulu et al., 2017). Interestingly, the SO-*m*AIEC can produce LA and ALA oxylipins, presumably from the LA and ALA in soybean oil (Figure 5). However, the oxylipins that were elevated in the bacterial cultures were not the same as those that were elevated in the mouse intestinal epithelial cells and some oxylipins were decreased in both Conv and GF mice on the soybean oil diet (e.g., 12,13-DiHOME and 12,13-Di-HODE). A similar complex pattern of certain types of oxylipins being increased while others are decreased in colon biopsies from UC patients has been reported previously (Diab et al., 2019). A causal role for oxylipins in IBD remains to be determined in both human and mouse models.

### Role of decreased endocannabinoids and EPA in SO-induced susceptibility to colitis

In addition to the LA/ALA oxylipins, the other most significantly dysregulated group of metabolites in both the bacterial and mouse metabolomic analyses were endocannabinoids: the presence of soybean oil caused a significant decrease in two of these compounds in both systems, EPEA and POEA (Figures 5 and Figure 7). Particularly intriguing was the fact that SO-*m*AIEC grown in the presence of soybean oil had lower levels of the endocannabinoids than the bacteria grown in the absence of soybean oil, suggesting that some component of the oil, or some pathway activated by soybean oil in the bacteria, resulted in decreased production and/or increased turnover of the endocannabinoids. One such compound is PEA which has been shown to increase the resistance to bacterial infections (Redlich et al., 2014). These results are consistent with decreased endocannabinoid levels being associated with increased colitis (Diab et al., 2019). Endocannabinoids (and cannabis) are increasingly being explored for potential therapeutic activity against IBD (Cani et al., 2016; Naftali, 2019).

In contrast to LA and ALA, EPA -- one of the best studied omega-3 PUFAs in terms of beneficial health effects -- was significantly decreased in both the *in vitro* and *in vivo* metabolomes in the presence of soybean oil. This is even though its parent fatty acid, ALA, is enriched roughly 30-fold in the soybean oil diet (3 kcal% in SO+f vs 0.1 kcal% in Viv chow) and suggests that some aspect of the ALA to EPA metabolic pathway in the host cells is compromised by dietary soybean oil. Additionally, we observed higher levels of EPA in GF compared to Conv mice, suggesting that the gut microbiota also influence EPA levels. This is consistent with the fact that EPA is decreased in SO-*m*AIEC *in vitro;* although it remains to be determined whether it is the SO-*m*AIEC that is causing the decrease of this beneficial fatty acid *in viv*o.

Many EPA metabolites, which are considered to be anti-inflammatory -- e.g., DiHETEs, HEPEs, PGF3a (Gabbs et al., 2015; Ramstedt et al., 1984) -- were also decreased by the SO+f diet (Supplementary Figure 5B, 7). The decrease was observed in both GF and Conv mice, suggesting a host rather than a bacterial effect. Additionally, the fact that some of these potentially beneficial oxylipin metabolites of EPA were increased in SO-*m*AIEC grown in media supplemented with soybean oil (e.g., PGF3a and 14,15-DiHETE, Supplementary Figure 5), suggests a complex interplay between the microbiome and the metabolome.

Overall, the decrease in the omega-3 EPA and several of its metabolites, coupled with the increase in the omega-6 LA and its metabolites, is expected to generate a pro-inflammatory state commonly found in IBD patients (Lee et al., 2018). Consistently, intestinal immune dysregulation was noted in WT mice fed soybean oil (Figure 2) and could be related to a pro-inflammatory state caused by an elevated omega-6/omega-3 ratio (Calder, 2005; Chapkin et al., 2007).

In conclusion, this study shows that a soybean oil-based diet similar to that currently consumed in the U.S. and increasingly prevalent worldwide can increase the susceptibility to colitis in rodents. The mechanism(s) thus far appears to involve dysregulation of the gut microbiome, potentially triggered by the high LA content of soybean oil, which could provide an intestinal pathobiont like SO-*m*AIEC a competitive growth advantage. This bacterium (SO-*m*AIEC) in turn contributes to an altered gut metabolome with increased oxylipins and decreased endocannabinoids and EPA. The net result is a proinflammatory environment characteristic of IBD. One or more of these conditions may disrupt the imbalance of the HNF4α isoforms, leading to impaired barrier function. Finally, two other high fat diets, olive oil and a low LA soybean oil (Plenish), did not show the same negative effects as the soybean oil diet indicating that the type of dietary fat can influence intestinal health. Follow-up studies are warranted to verify that the effects observed in mice also occur in humans.

## METHODS

### Animals and diets

Care and treatment of animals was in accordance with guidelines from and approved by the University of California, Riverside Institutional Animal Care and Use Committee and followed NIH guidelines. Young adult male mice were maintained on a 12:12 hour light-dark cycle in either a conventional, non-specific-pathogen free vivarium or in a gnotobiotic facility, as indicated. Conventionally raised, wild-type (WT) C57BL/6N (Charles River), exon swap HNF4α (α1HMZ) (Briancon and Weiss, 2006) and IL-10^-/-^ mice (Kühn et al., 1993) (Jackson Labs, Stock#: 002251) were used. The IL-10^-/-^ mice were in the B6 strain, which is known to develop the milder form of spontaneous colitis (JAX Labs, catalog # 002251). Germ-free mice (C57BL/6N, Taconic) were raised under gnotobiotic conditions, as described previously (Alavi et al., 2020). All the mice were fed a standard vivarium chow from Newco until they were weaned 21 days after birth (autoclaved LabDiet 5K52 for the germ-free mice and non-autoclaved LabDiet 5001 for the conventionally raised mice). Post weaning, the mice were either continued on the low-fat vivarium chow or fed one of four isocaloric high fat diets for up to 24 weeks: a soybean oil-based high fat diet with (SO+f) or without added fiber (SO), a low LA soybean oil diet (Plenish) or an olive oil diet. All four high fat diets have 35 kcal% fat and were formulated by Research Diets Inc. See Supplementary Table 1 for detailed composition of diets. All mice had *ad libitum* access to food and water. At the end of the study, mice were euthanized by carbon dioxide inhalation, in accordance with NIH guidelines.

### Dextran sulfate sodium (DSS) treatment

WT or α1HMZ mice that had been on the diets for 8 to 15 weeks were treated with 2.5% dextran sodium sulfate salt (DSS) (reagent grade, MW 3.6–5 kDa, MP Biomedicals, #160110, Santa Ana, CA) in water *ad libitum* for six days and sacrificed immediately or allowed to recover up to 3 days with tap water. Mice were continued on the same diet before, during and after DSS treatment and monitored daily for changes in body weight, stool consistency, ruffled fur and activity level. Presence of blood in the stool was checked every other day using Hemoccult Dispensapak Plus (catalog no. 61130, Beckman Coulter).

### Disease activity index

A composite disease activity index (DAI) score was calculated based on the following criteria (Lo Sasso et al., 2020; Montbarbon et al., 2013): 1) weight loss 2) colonic length to weight ratio 3) hemoccult reading 4) gross morphological changes in colon. See Supplementary Table 2 for details on scoring criteria.

### Linoleic Acid Gavage

WT mice on vivarium chow were gavaged with either pharma grade linoleic acid (catalog no. 39269-10G, Millipore Sigma) or water for 3 days using disposable animal feeding needles (catalog no. 01-208-87, Thermo FisherScientific) and syringes (catalog no. 14-823-434, BD Slip tip, Thermo FisherScientific). Daily dose of LA gavage was 0.26 mg/kg body weight per day; together with the LA in the Viv chow this is similar to the amount the mice ingested on the soybean oil-enriched diet (i.e., 18.6 kcal% LA) and is similar to what has been used in previous studies (Rodrigues et al., 2012). Fecal samples were collected and stored at −80°C prior to the first gavage and at the end of the experiment (i.e., 24 hours after the third gavage). These samples were used for bacterial rRNA internal transcribed spacer (ITS) library construction.

### Tissue collection

Tissues were collected and snap-frozen in liquid nitrogen prior to storage at −80°C for immunoblotting and metabolomic analysis or fixed in 10% neutral-buffered formalin for 24 hours before storing in 30% sucrose plus PBS solution at 4°C for subsequent histological analysis. Liver and adipose tissue (mesenteric, perirenal, gonadal and flank subcutaneous) were excised and weighed.

### Tissue Embedding, Sectioning and Staining

The entire large intestine was excised, and its length measured. A 1.0 to 1.5-cm piece adjacent to the rectum was cut and fixed in 10% neutral-buffered formalin as above, embedded in optimal cooling temperature (OCT) compound and sectioned at 5-µm thickness on a Microm Cryostat and stored at −20°C. All slides were rehydrated in 95% ethanol for 7 minutes, tap water for 7 minutes, ddH20 for 2 minutes, and stained in hematoxylin (Ricca Chemical) for 40 seconds. Slides were then dipped in tap water for 30 seconds, running tap water for 90 seconds, 95% ethanol for 15 seconds and subsequently counterstained in eosin (Sigma-Aldrich) for 3 seconds and dipped in 95% and 100% ethanol two and three times, respectively, for 20 seconds each time. Slides were left in Citrisolve (Fisher Scientific) for at least 40 seconds. This staining process was completed in succession in a single session. Slides were fixed and preserved with Permount (Fisher Chemicals). Histology images were captured on an Evos Microscope (Life Technologies). Crypt length and submucosal thickness were measured using SPOT Imaging software (Sterling, MI).

### Immunoblot Analysis

Whole cell extracts were prepared from tissues stored in liquid nitrogen and analyzed by immuno-chemiluminescence after determination of protein concentration by the Bradford Assay, as described previously (Chellappa et al., 2016; Maeda et al., 2006). The protein extracts were separated by 10% sodium dodecyl sulfate-polyacrylamide gel electrophoresis (SDS-PAGE) and transferred to Immobilon membrane (EMD Millipore, Billerica, MA). The membrane was blocked with 5% nonfat milk for 30 minutes, incubated in primary antibodies (mouse monoclonal anti-HNF4α P1 and P2; catalog no. PP-K9218-00 and PP-H6939-00, respectively, R&D Systems) in 1% milk, overnight at 4°C. After several washes in TBST (Tris-buffered saline, 0.1% Tween 20), the blots were incubated in horseradish peroxidase (HRP)-conjugated goat anti-mouse (GαM-HRP) secondary antibody (Jackson ImmunoResearch Laboratories) for 40 minutes at room temperature followed by three 5-minutes washes in TBST and two 5-minute washes in TBS (Tris-buffered saline). Blots were developed using SuperSignal™ West Pico PLUS Chemiluminescent Substrate (Thermofisher Scientific) and imaged in a Chemi-Doc imaging system (Bio-Rad). Coomassie staining of blots verified equal protein loading. The blots were re-probed for beta-actin (rabbit anti-actin; catalog no. A2066, Sigma) by washing twice in TBST before incubating in a stripping buffer (0.5 M NaOH solution) at room temperature for 5 minutes with shaking, washed twice with TBST and once with TBS (3 minutes each). Blots were blocked in 5% milk and the immunoblot procedure described above was followed.

### In-vivo Permeability Assay

Mice were fasted overnight on wood chip bedding (Newco Specialty, catalog # 91100). After 15 hours, mice were weighed and gavaged with FITC-Dextran (FD-4 Sigma) diluted in water at a dose of 600 µg/gm body weight. The gavage was performed under yellow lights and staggered 2 minutes between animals. Mice were sacrificed 4 hours after gavage and blood was collected via cardiac puncture (BD 3-ml Luer-Lok Syringes, catalog no. 14-823-435, Fisher and BD 26 G ⅝ inch hypodermic needles, catalog no. 14-826-6A, Fisher) and transferred to 1.5-ml Eppendorf tubes. Samples were placed on ice for 45 minutes and then centrifuged at 9.3 rcf for 15 minutes after which serum was collected into a fresh tube. Samples were loaded into black 96-well plates (Corning, catalog no. 3991) in triplicate, at a dilution of 1:5 in water. Serum FITC-Dextran concentration was determined on a Veritas Microplate Luminometer (Turner Biosystems, Sunnyvale, CA), GloMax software (Promega, Madison, WI), using excitation/emission wavelengths of 490/520 nm(Shawki et al., 2020). The relative fluorescence units obtained for the samples were compared to the values obtained from a standard curve generated by diluting the fluorophore stock in water.

### Intestinal immune cell profile

#### IEL and LPL isolation

Intraepithelial (IELs) and lamina propria lymphocytes (LPLs) were isolated from mouse small intestines as described previously (Couter and Surana, 2016). Briefly, the entire small intestine was excised and gently flushed with cold PBS and mesenteric fat and Peyer’s Patches were dissected away. The intestine was cut into 3 to 4-inch segments; each segment was rolled on a paper towel moistened with Gibco RPMI 1640 Media to remove any residual fat tissue. The segments were inverted using curved forceps and placed in 30 ml extraction medium -- RPMI plus 93 µl 5% (w/v) dithiothreitol (DTT), 60 µl 0.5 M EDTA and 500 µl fetal bovine serum per small intestine, ∼40 cm long -- and stirred at 500 rpm for 15 minutes at 37°C. A steel strainer was used to separate tissue pieces from the IEL-rich supernatant, which was placed on ice. Residual mucus from the tissue segments was removed by blotting on a dry paper towel. These fragments were then put in a 1.5-ml Eppendorf tube with 600 µl of digestion medium: 25 ml RPMI plus 12.5 mg dispase, 37.5 mg collagenase II and 300 µl FBS. Dispase (Gibco, catalog no. 17105041) and collagenase (Gibco, catalog no. 17101015) were added immediately before use. The tissue was minced inside the tube using scissors, put in a cup containing 25 ml of digestion media and stirred at 500 rpm for 15 minutes. Any large chunks of tissue were broken up by pipetting up and down with a serological pipette and stirring at 37°C was continued for an additional 15 minutes. Digested tissue and the IEL containing supernatant were passed through a 100-µm cell strainer into a 50-ml tube followed by a rinse with 20 ml of RPMI containing 10% FBS. The filtered solution was centrifuged at 500 x g for 10 minutes at 4°C; the supernatant was carefully decanted, and the pellet resuspended in 1 ml of RPMI containing 10% FBS. The resuspended cells were filtered through a 40-µm cell strainer into a 50-ml tube followed by a rinse with 20 ml of RPMI containing 10% FBS. The filtered solution was centrifuged at 500 x g for 10 minutes at 4°C; the supernatant was carefully decanted, and the pellet resuspended in 1 ml of RPMI containing 2% FBS. This suspension was then used for flow cytometry as described below.

#### PBMC isolation

Mice were euthanized and blood was collected and immediately mixed with 1 ml 4% sodium citrate followed by addition of Histopaque-1077 (catalog no:10771, Sigma-Aldrich). This mix was centrifuged at 400 g for 30 minutes at room temperature. PBMCs were carefully aspirated from the interphase and washed with culture medium.

#### Flow cytometry

Isolated PBMC, IELs and LPLs were washed in fluorescence activated cell sorting buffer (FACS) buffer, incubated with Fc block (25 µg/mL αCD16/32 and 10 µg/ml rat IgG) and stained for 30 minutes with flow antibodies: F4/80 (Cl:A3-1, Bio Rad MCA497FB), SiglecF (E50-2440, BD Biosciences 562757), CD4 (RM4-5, BD Biosciences 550954), Ly6C (HK 1.4, Biolegend 128018), CD11b (M1/70, Biolegend 101226), CD11c (N418, Biolegend 117310), Ly6G (1A8, Biolegend 127628), MHCII (M5/114.15.2, Biolegend 107622), CD19 (1D3, BD Biosciences 562956), CD115 (AFS98, Biolegend 135517) and CD8 (53-6.7, Biolegend 100742). All cells were acquired on the BD LSRII (BD Biosciences) and analyzed on FlowJo (FlowJo^TM^ v10). Cell populations were identified as: macrophage (CD11b^+^ F4/80^+^), eosinophils (CD11b^+^ SiglecF^+^), monocytes (CD11b^+^ Ly6C^+^), neutrophils (CD11b^+^ Ly6G^+^), CD4^+^ T cells (CD4^+^ CD8^-^), CD8^+^ T cells (CD8^+^ CD4^-^) and B cells (CD4^-^ CD19^+^).

### Isolation of intestinal epithelial cells (IECs) for metabolome and microbiome analysis

Intestinal epithelial cells (IECs) were isolated for microbiome and metabolomic analysis as previously described (Shawki et al. 2020). Bacterial DNA was isolated from the IECs using the DNeasy PowerSoil Kit (Qiagen, Valencia, CA) with a 30-second bead-beating step using a Mini-Beadbeater-16 (BioSpec, Bartlesville, OK). For metabolomics analysis, the cells were flash frozen in liquid nitrogen and stored at −80°C until processing.

### Bacterial rRNA internal transcribed spacer (ITS) library construction and sequencing

Illumina bacterial rRNA ITS gene libraries were constructed as follows: PCR was performed in an MJ Research PTC-200 thermal cycler (Bio-Rad Inc., Hercules, CA) in 25-μl reactions containing 50 mM Tris (pH 8.3), bovine serum albumin (BSA) at 500 μg/ml, 2.5 mM MgCl_2_, 250 μM of each deoxynucleotide triphosphate (dNTP), 400 nM of the forward PCR primer, 200 nM of each reverse PCR primer, 2.5 μl of DNA template and 0.625 units JumpStart Taq DNA polymerase (Sigma-Aldrich, St. Louis, MO). PCR primers targeted a portion of the small-subunit (ITS-1507F,GGTGAAGTCGTAACAAGGTA) and large-subunit (ITS-23SR, GGGTTBCCCCATTCRG) rRNA genes and the hypervariable ITS region (Ruegger et al., 2014), with the reverse primers including a 12-bp barcode and both primers including the sequences needed for Illumina cluster formation; primer binding sites were the reverse and complement of the commonly used small-subunit rRNA gene primer 1492R (Frank et al., 2008) and the large-subunit rRNA gene primer 129F (Hunt et al., 2006). PCR primers were only frozen and thawed once. Thermal cycling parameters were 94°C for 5 minutes; 35 cycles of 94°C for 20 seconds, 56°C for 20 seconds, and 72°C for 40 seconds; followed by 72°C for 10 minutes. PCR products were purified using a Qiagen QIAquick PCR Purification Kit (Qiagen) according to the manufacturer’s instructions. DNA sequencing (single-end 150 base) was performed using an Illumina MiSeq (Illumina, Inc., San Diego, CA).

### Bacterial rRNA ITS sequence processing and analysis

The UPARSE pipeline was used for de-multiplexing, length trimming, quality filtering and amplicon sequence variant (ASV) picking using default parameters or recommended guidelines (Edgar, 2013) updated at http://www.drive5.com/usearch/manual10/uparse_pipeline.html. Briefly, after demultiplexing and using the recommended 1.0 expected error threshold, sequences were trimmed to a uniform length of 145 bp and then de-replicated. De-replicated sequences were subjected to error correction (denoised) and chimera filtering to generate zero radius operational taxonomic units (ZOTUs) using UNOISE3 (Edgar, 2016). An ASV table was generated using the otutab command. ASVs having non-bacterial DNA were identified by performing a local BLAST search (Altschul et al., 1990) of their seed sequences against the nucleotide database. ASVs were removed if any of the highest scoring BLAST hits contained taxonomic IDs within the rodent family, the kingdoms Fungi or Viridiplantae kingdoms, or PhiX. Taxonomic assignments of the bacterial ASVs were made by finding the lowest common taxonomic level of the highest BLAST hits excluding unclassified designations. Data were normalized within each sample by dividing the number of reads in each ASV by the total number of reads in that sample. The bacterial rRNA ITS sequences have been deposited in the National Center for Biotechnology Information (NCBI)’s Sequence Read Archive (SRA) under the SRA BioProject Accession PRJNA622821.

#### Bacterial rRNA ITS sequence analysis

QIIME (Caporaso et al., 2010) was used to create the abundance tables for the phylotypes at various taxonomic levels. Correlation analyses and plots, as well as bacteria species plots, were performed using Prism (GraphPad, La Jolla, CA).

#### Beta Diversity Plot

QIIME (Caporaso et al., 2010) was used to calculate a Hellinger beta diversity distance matrix (for the LA gavage experiment), which was depicted using principal coordinates analysis and statistically assessed using Adonis PERMANOVA tests.

### Isolation of *m*AIEC

The *m*AIEC strain used in this study (UCR-SoS5, referred to as SO *m*AIEC in the Results) was isolated from subcutaneous fat collected from mice fed the high soybean oil diet with fiber (SO+f) using a selective medium, *E. coli* ChromoSelect Agar B, as described by the manufacturer (Sigma-Aldrich, St. Louis, MO). The strain was purified by selecting a colony from the selective media and then performing two successive streak plating procedures on LB agar to obtain single colonies. The strain was confirmed to have the identical rRNA ITS nucleotide sequence as the *E. coli* phylotype identified by the Illumina sequence analysis by PCR amplifying the rRNA ITS region of the strain and sequencing the amplicons using the Sanger method.

### *m*AIEC phenotypic tests

Adherence and invasion ability of *m*AIEC to Caco-2 brush border epithelial (Caco2-BBe) cells was examined using previously described methods (Shawki et al. 2020). Intracellular replication of *m*AIEC was performed in J774A.1 cells (ATCC TIB-67), murine macrophages maintained in RPMI plus 10% FBS, penicillin (100 U/ml), and streptomycin (100 µg/ml) until the day before the experiment, at which point they were maintained in the same media without antibiotics. Intracellular replication of *m*AIEC in J774A.1 cells was determined by a gentamicin survival assay using 24-well plates with 2 ml of media per well. J774A.1 monolayers were infected at an MOI of 20 bacteria/macrophage and incubated for 2 hours at 37°C with 5% CO2. Infected macrophages were washed twice with PBS and incubated 1 hour in RPMI plus 10% FBS and 150 mg/ml gentamicin to kill extracellular bacteria. The J774A.1 cells were washed once with PBS and then lysed by incubation with 1 ml of 1% Triton-X 100 for 30 minutes; using two extra uninoculated wells, an average J774A.1 cell count per well was obtained at this point. The lysed cell solutions (1 ml) were collected, centrifuged at 14,000 x g for 10 seconds, the supernatants were decanted, and the cells were resuspended in 120 µl PBS. The lysed cell solutions were serially diluted, spread-plated on LB agar, incubated overnight at 37 °C, and the bacteria were enumerated. This process was repeated for another plate but after the 1-hour 150 mg/ml gentamycin incubation described above, the cells were washed once with PBS, and then RPMI plus 10% FBS and 20 mg/ml gentamicin was added and the cells were allowed to incubate for another 23 hours at 37°C. Intracellular replication of the *m*AIEC and control bacteria was expressed as the mean percentage of the number of bacteria recovered 24 hours post-infection divided by the number of bacteria one hour post-infection, with the two J774A.1 cell counts being used to normalize the bacterial counts. Control bacteria were the LF82 human AIEC (kindly provided by the late Dr. Arlette Darfeuille-Michaud) and K12 (a noninvasive *E. coli*, ATCC 25404).

### Culture of *m*AIEC for metabolomic analysis

The *m*AIEC was grown aerobically in LB broth with 10% soybean oil for 20 hours at 37°C with shaking at 350 rpm. The bacteria were collected by centrifugation at 14,000 x g for 30 seconds and the supernatant removed; this procedure was repeated two more times to remove as much of the soybean oil as possible. The bacterial pellets were flash-frozen in liquid nitrogen and stored at −80°C until they were analyzed. Controls were the *m*AIEC grown in LB broth without soybean oil as well as uninoculated LB with and without soybean oil that was subjected to incubation for 20 hours at 37°C with shaking at 350 rpm.

### Sole carbon source experiments

*m*AIEC or *E. coli* K-12 were grown aerobically in LB media for 20 hours at 37°C at 300 rpm and then diluted (1:50) in minimum essential medium (standard M9 medium but without glucose, supplemented with vitamins (ATCC, catalog no. MD-VS) and trace minerals (ATCC, catalog no. MD-TMS). This medium was supplemented with either 2 mM linoleic acid dissolved in ethanol (Cayman Chemicals, catalog no. 9015050) or an equal volume of ethanol. A third treatment was included for *E. coli* K-12 in which the M9 medium was supplemented with 0.4% glucose. The OD600 measurements were taken on a Synergy HTX Microplate reader at 0, 30, 90, 150, 180 and 240 minutes after inoculation. To show viability of the K-12 bacteria after LA and ethanol treatment, glucose was added at a concentration of 0.4% after 240 minutes and two additional OD readings, at 360 and 1440 minutes were taken. There were three replicate cultures per condition: results from one representative experiment out of at least two independent experiments are shown.

### Metabolomic Analysis

#### Cell pellet: non-esterified oxylipins, endocannabinoids and polyunsaturated fatty acids extraction

Cell pellets (of IECs and bacteria) were weighed prior to extraction. Oxylipins, endocannabinoids and polyunsaturated fatty acids (PUFAs) were isolated by liquid extraction protocol using acetonitrile/isopropanol/water mixture [3:3:2 v/v] from approximately 80 mg of cell pellet and quantified by UPLC-MS/MS using internal standard methods. Briefly, ∼80 mg of cell pellet was mixed with 20 µL BHT/EDTA (1:1 MeOH:water), 20 µL of 1250 nM deuterated oxylipins and endocannabinoids surrogates in methanol and 20 µL of 1-cyclohexyl ureido, 3-dodecanoic acid (CUDA) and 1-phenyl ureido 3-hexanoic acid (PUHA) at 5 µM in 1:1 methanol:acetonitrile. Samples were then homogenized using Geno/Grinder 2010 after addition of 0.5 mL of acetonitrile/isopropanol/water (3:3:2), together with three 3-mm stainless steel beads. Homogenate was centrifuged at 15000 rcf for 10 minutes and filtered through a 0.1 µm PVDF spin filter and collected for mass spectrometry analysis described below.

#### Bacterial culture media: non-esterified oxylipins, endocannabinoids and polyunsaturated fatty acids extraction

Non-esterified oxylipins, endocannabinoids and polyunsaturated fatty acids were isolated using solid phase extraction with 60 mg Hydrophilic-Lipophilic-Balanced columns (Oasis, Waters Corporation, Milford, MA). Briefly, columns were washed with one column volume of ethyl acetate followed by two column volumes of methanol and further conditioned with two column volume of 5% methanol, 0.1% acetic acid in water. Next, columns were spiked with 5 µL BHT/EDTA (1:1 MeOH:water) and 5 µL of 250 nM deuterated oxylipins and endocannabinoid surrogates in methanol. Samples (200 µL) were mixed with 800 µL of 5% methanol, 0.1% acetic acid in water, transferred onto the column and extracted under the gravity. Columns were washed with 1 column volume of 30% methanol, 0.1% acetic acid in water. Analytical targets were eluted with 0.5 mL methanol followed by 1.5 mL ethyl acetate. Eluents were dried under the vacuum, reconstituted in 50 µL of 1-cyclohexyl ureido, 3-dodecanoic acid (CUDA) and 1-phenyl ureido 3-hexanoic acid (PUHA) at 5 µM in 1:1 methanol:acetonitrile, filtered through 0.1 µm PVDF spin filter and collected for mass spectrometry analysis described below.

#### Mass spectrometry analysis

Residues in extracts were separated on a 2.1 mm x 150 mm, 1.7 µm BEH C18 column (Waters, Milford, MA) and detected by electrospray ionization with multi reaction monitoring on a API 6500 QTRAP (Sciex; Redwood City, CA) and quantified against 7-9 point calibration curves of authentic standards using modifications of previously reported methods (Agrawal et al., 2017).

### Statistical analysis

Data are presented as the mean ± standard error of the mean (SEM) using GraphPad Prism 6. One-way ANOVA or repeated measures (RM) ANOVA followed by post-hoc testing (Tukey’s for multiple comparisons or Sidak for pairwise comparisons, as specified in the figure legends) or Student’s T-test were used as appropriate and are indicated in the figure legends. Statistical significance was set at an alpha level of 0.05 with a P*-*value less than or equal to 0.05 being considered significant. Linear regression analysis was performed between body weight, adipose tissue weight, liver as percent body weight, colon length and relative abundance of *m*AIEC in intestinal epithelial cells. The following cut-offs were used to determine significance: Pearson’s coefficient r > 0.5 with P ≤ 0.05. For the *metabolomics data,* outliers were first removed using the robust Huber M test and missing data were imputed using multivariate normal imputation. Further, variables were clustered separately for each cell type using the JMP variable clustering algorithm, an implementation of the SAS VARCLUS procedure, and converted into cluster components for data reduction. Curated metabolomics data are presented in Supplementary Table 4. Cluster components were used for PCA analysis of experimental samples to provide an overview of metabolic changes.

