## Supplementary Figures for "Diet High in Soybean Oil Increases Susceptibility to Colitis in Mice"

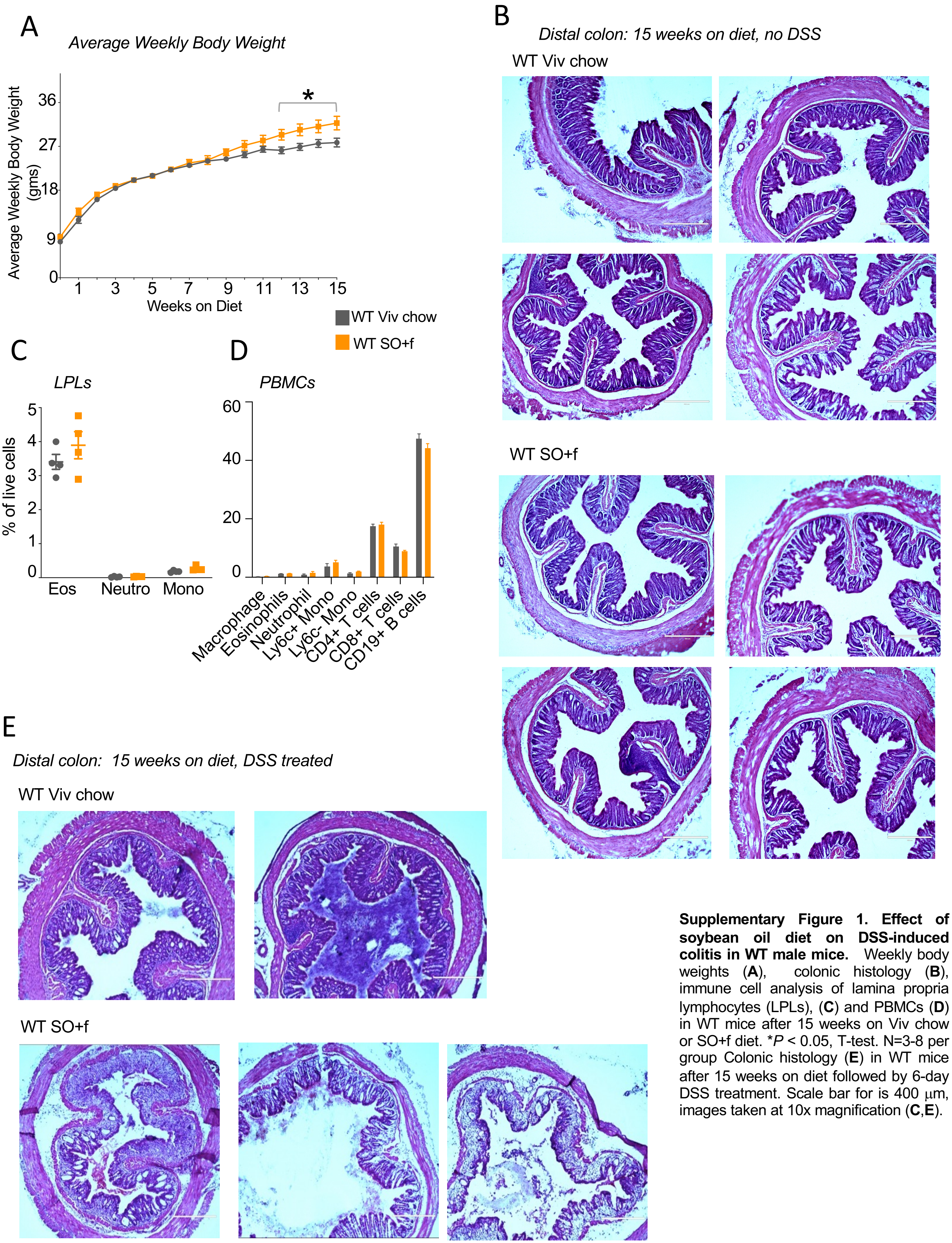


**Supplementary Figure 1. Effect of soybean oil diet on DSS-induced colitis in WT male mice.** Weekly body weights (**A**), colonic histology (**B**), immune cell analysis of lamina propria lymphocytes (LPLs), (**C**) and PBMCs (**D**) in WT mice after 15 weeks on Viv chow or SO+f diet. *$P < 0.05$, T-test. N=3-8 per group Colonic histology (**E**) in WT mice after 15 weeks on diet followed by 6-day DSS treatment. Scale bar for is 400 μm, images taken at 10x magnification (**C**,**E**).

A

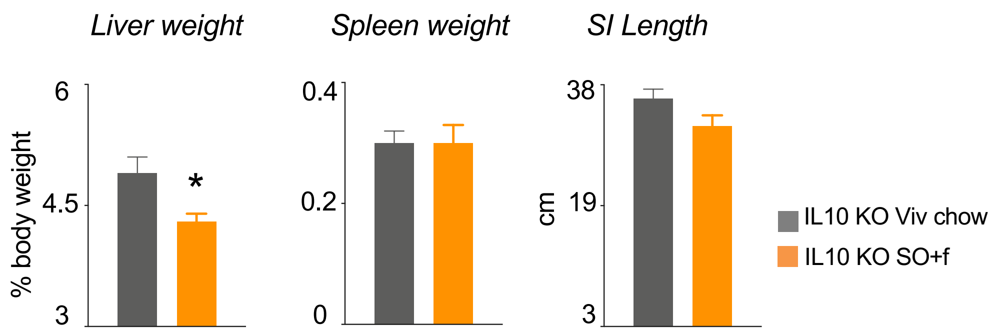


B

*Distal colon: 10 weeks on diet*

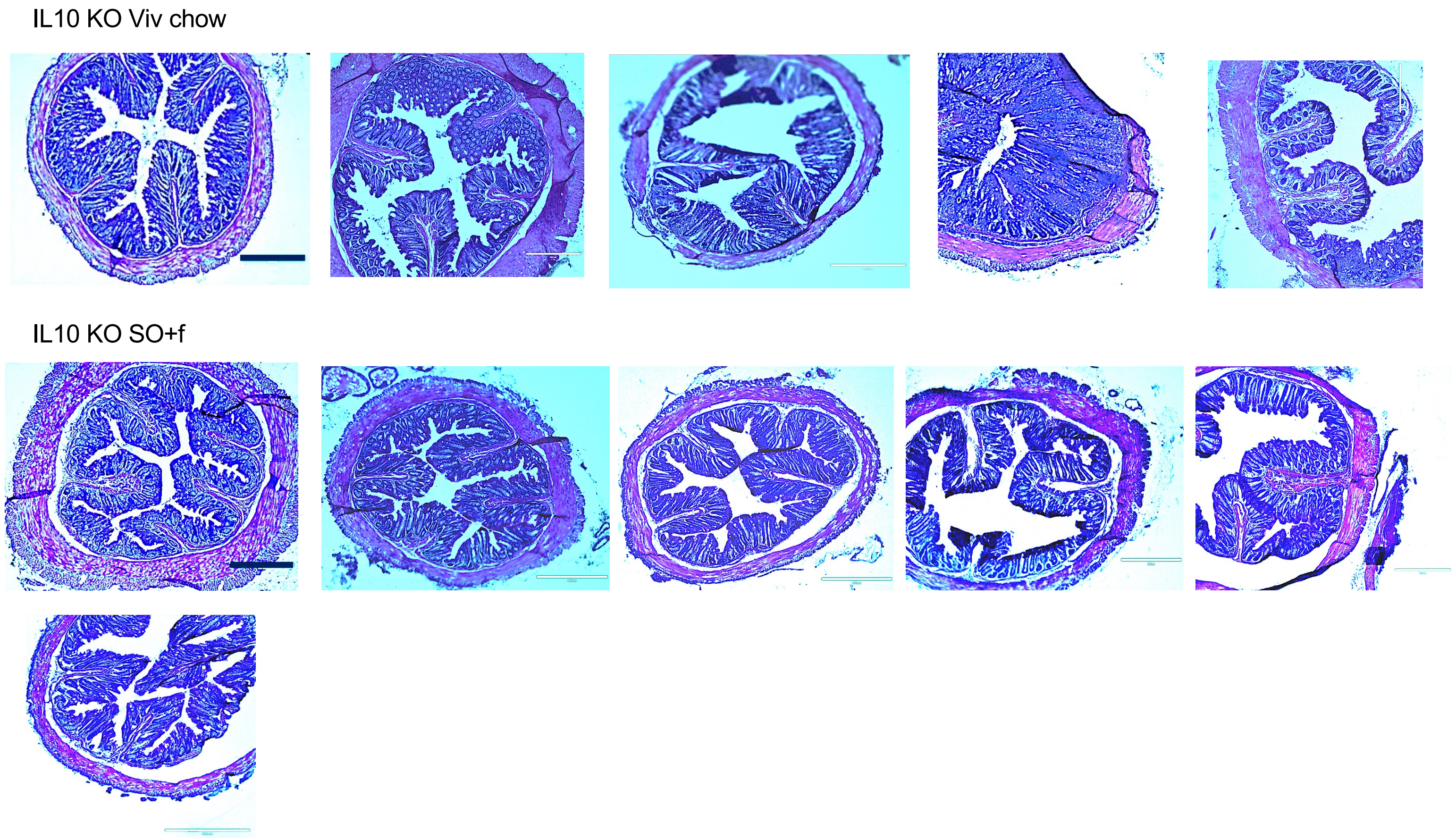


**Supplementary Figure 2. Effect of soybean oil diet on IL10 deficient male mice.**
Liver and spleen weights as percent of body weight and small intestinal length (**A**) and colonic histology (**B**) in IL10 KO mice after 10 weeks on Viv chow and SO+f diet. Scale bar for is 400 μm, images taken at 10x magnification. * *P* < 0.05 vs other diet, T-test. N=5-12 per group.

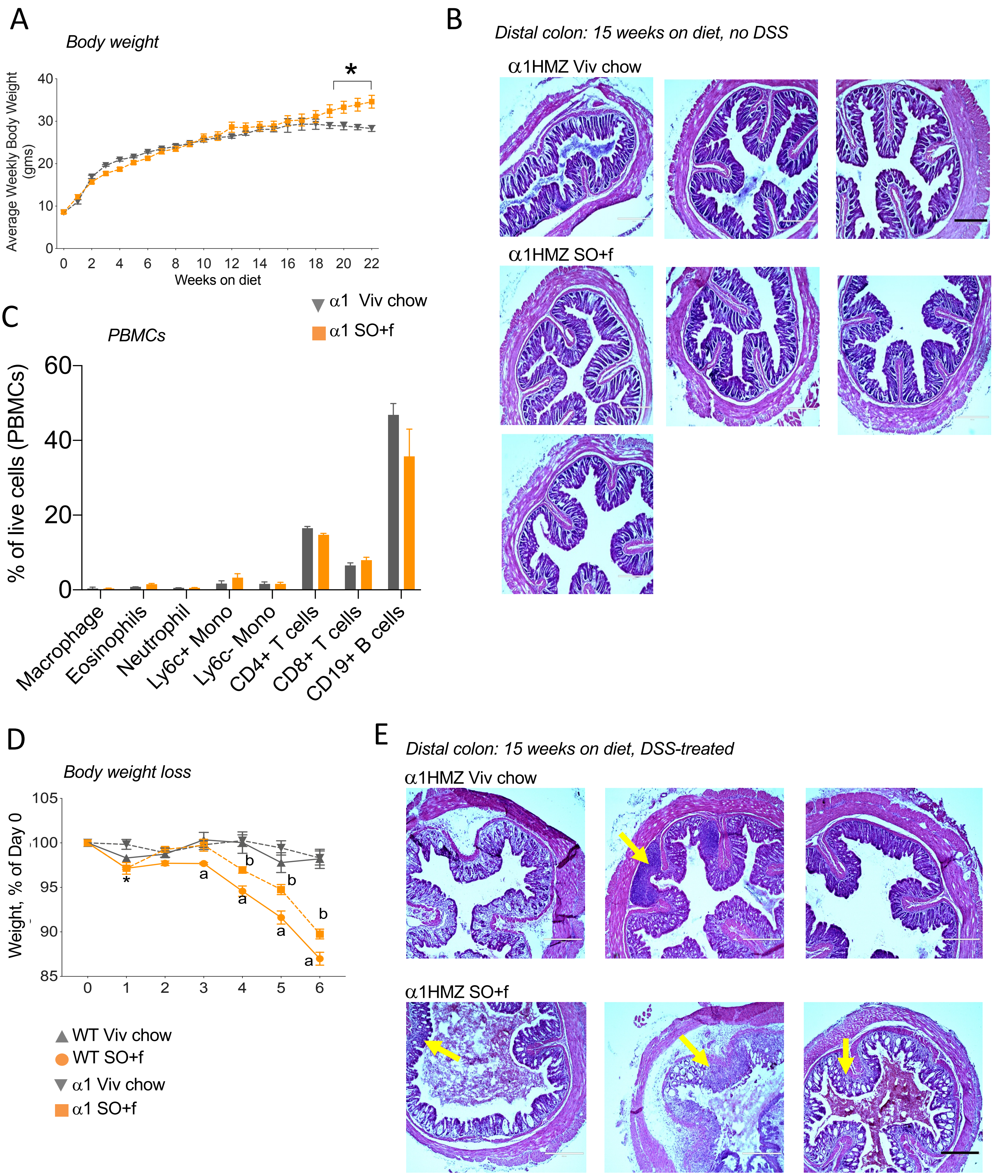


**Supplementary Figure 3. Effect of soybean oil diet on DSS-induced colitis in α1HMZ male mice.**
Weekly body weights up to 22 weeks on diet (**A**), colonic histology (**B**) and immune analysis in peripheral blood mononuclear cells (PBMCs) after 15 weeks on diet in α1HMZ mice (**C**). **D)** Comparison of % body weight loss between WT and α1HMZ mice on Viv chow or SO+f for 15 weeks, treated with 2.5% DSS in drinking water for 6 days (same data as in Fig. 1A and 2A). [*] α1HMZ SO+f vs α1HMZ Viv, [a] WT SO+f vs WT Viv, [b] α1HMZ SO+f vs WT SO+f: $P < 0.05$ vs other diet, repeated measures 2-way ANOVA, Tukey's post-hoc. N=3-4 per group **E)** Colonic histology in α1HMZ mice after 15 weeks on diet followed by 6-day DSS treatment. Arrow: immune cell infiltrate.

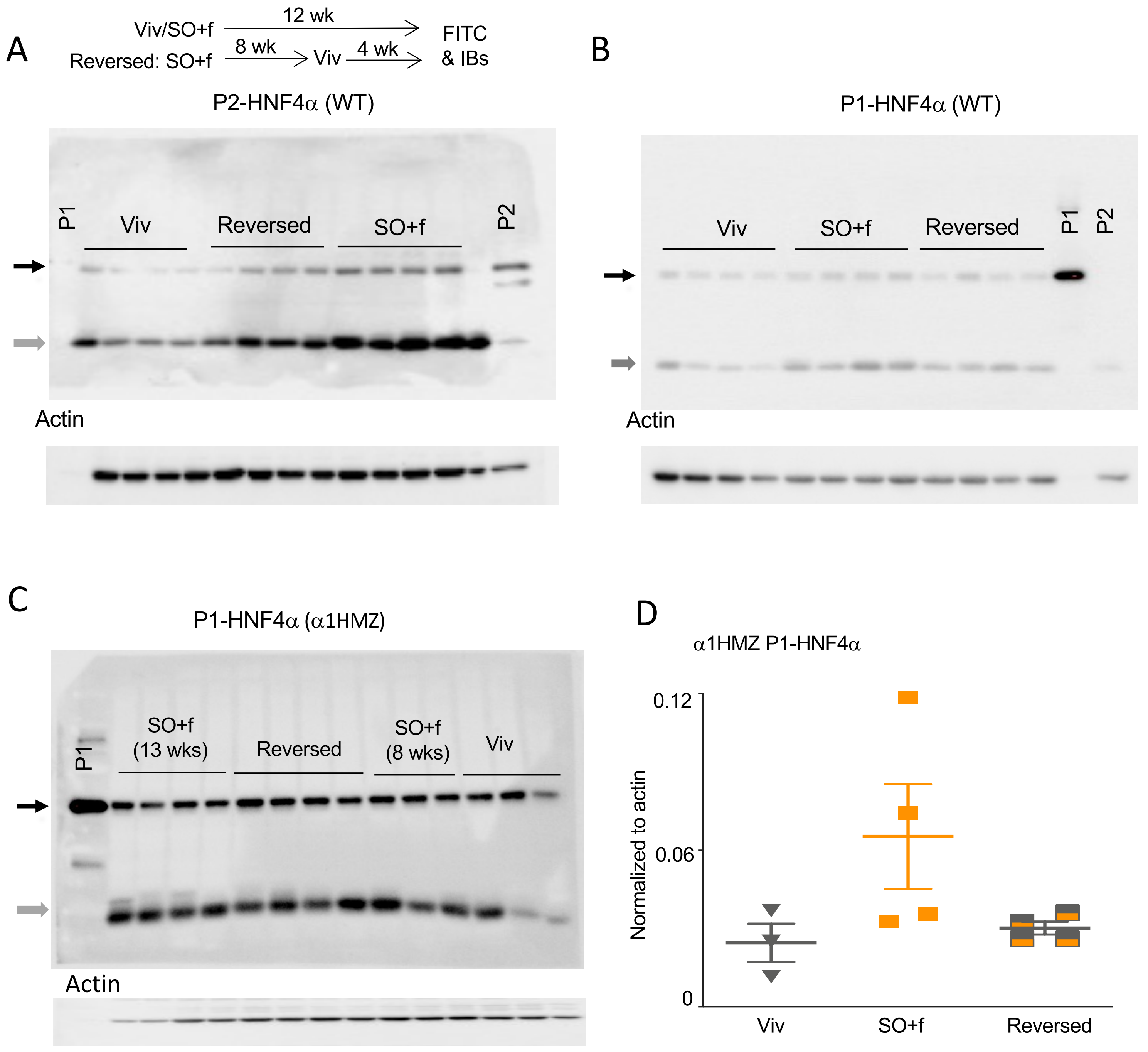


**Supplementary Figure 4. Effect of soybean oil diet on HNF4α isoform balance in the colon. A-C)** HNF4α immunoblots of whole cell extracts (WCE, 30μg) from distal colon of mice fed Viv chow or SO+f diet for 12 weeks or SO+f for 8 weeks followed by Viv chow for 4 weeks (reversed) (quantification shown in Figure 3F)**.** Each lane contains WCE from a different mouse. P1 control-nuclear extract from HCT116 cells expressing P1-HNF4α ; P2 control-nuclear extract from α7HMZ mouse. Black arrow: HNF4α; Gray arrow: non-specific band. **D)** Quantification of the P1-HNF4α signal normalized to total protein, as determined by actin staining of the same blot. N=3-4 per group.

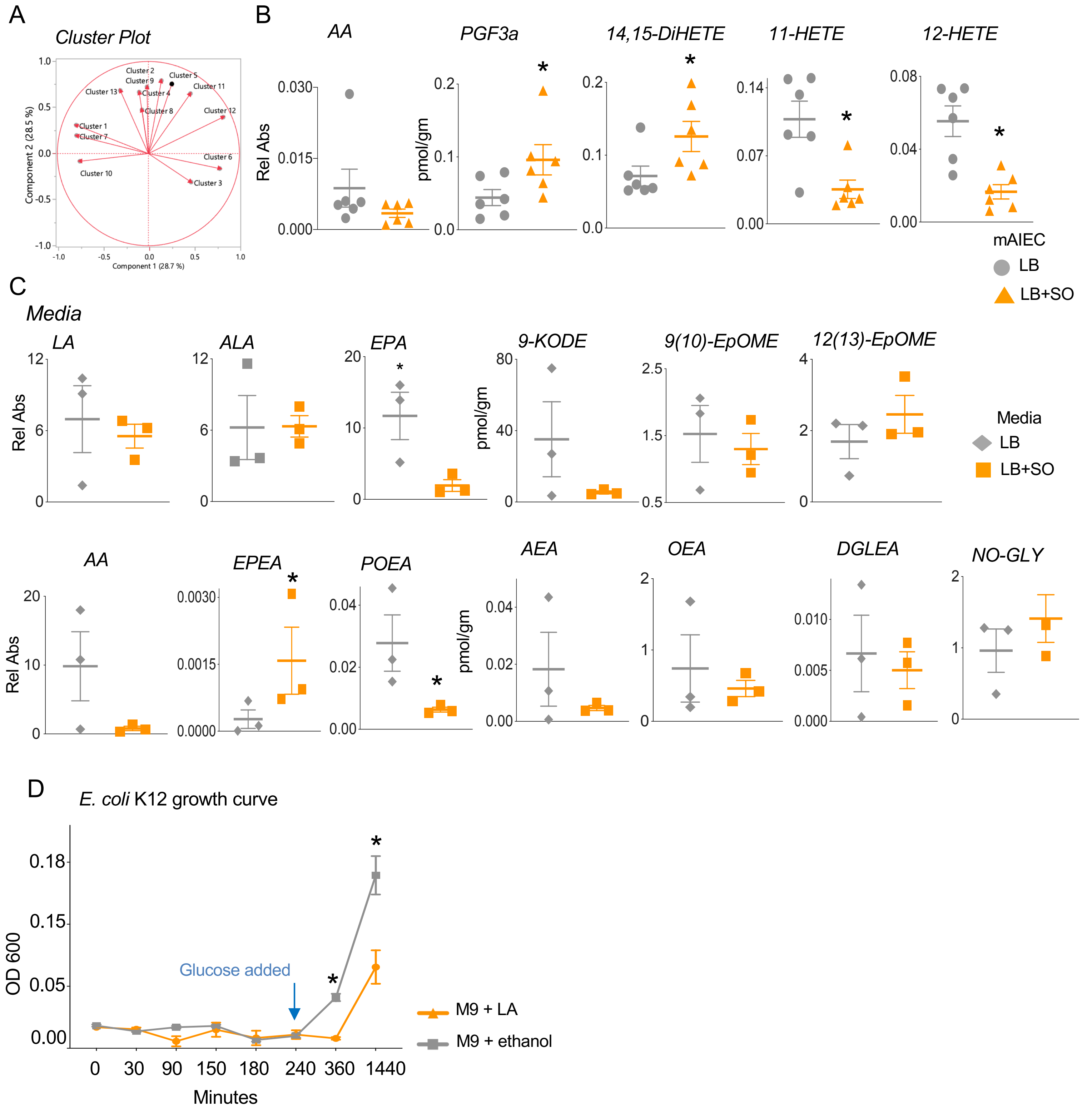


**Supplementary Figure 5. Effect of soybean oil on metabolome of *m*AIEC grown *in vitro* and of LA on growth curve of *E.coli* K12.** Cluster plot for Principal Components Analysis (PCA) shown in Figure 5A (see Supplementary Table 3 for details) (**A**) and absolute levels of fatty acids, oxylipin and endocannabinoid metabolites in *m*AIEC grown with or without soybean oil (SO) in media (**B**) and the corresponding media (Luria Broth, with or without added SO) (**C**). * $P < 0.05$, T-test. N=6-8 per group. **D)** Growth curves for the *E. coli* K12 grown in Minimum Essential Medium (M9) with LA or ethanol as the carbon source. * $P < 0.05$, T-test. N=3 cultures per group.

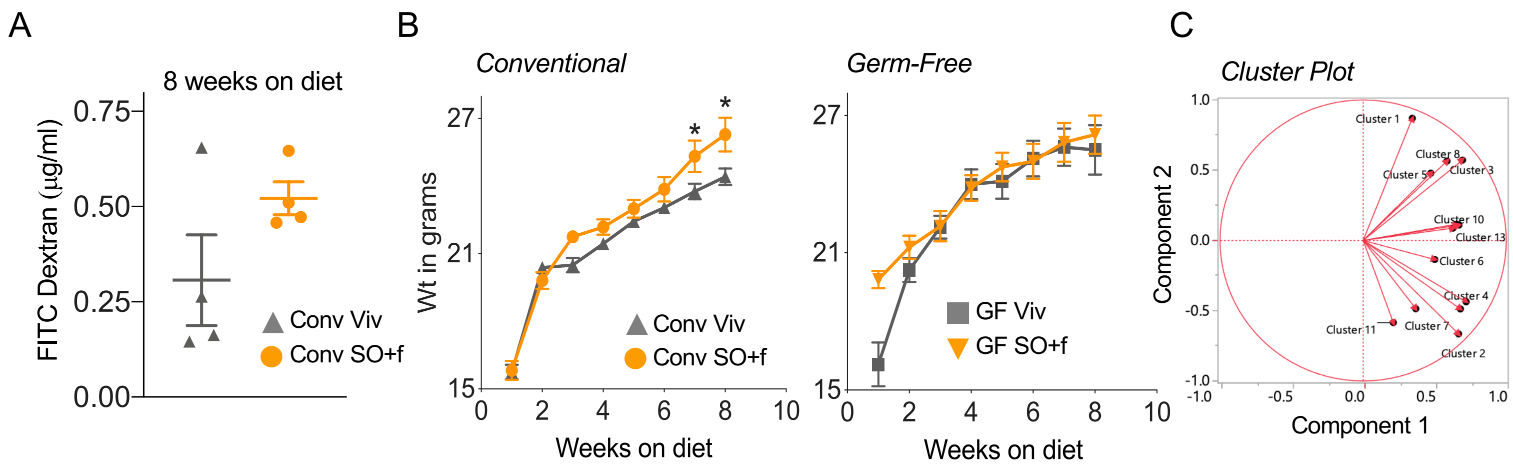


**Supplementary Figure 6. Effect of soybean oil on barrier function, body weight and metabolome of germ-free (GF) vs conventional (Conv) mice. (A)** Epithelial permeability in conventionally raised WT mice after 8 weeks on indicated diet**. B)** Weekly body weights in indicated groups of mice. * *P* < 0.05, T-test. N=6-8 per group. **C)** Cluster plot for PCA shown in Figure 6B of oxylipin and endocannabinoid metabolites in IECs from Conv or GF WT mice on SO+f or Viv diets for 8 wks. All mice are C57BL6/J strain adult males. N=6 per group.

| Dysregulated Metabolites in SO+f vs Viv chow in Conv mice only |
|---|
| 13-HODE |
| 13-HOTE |
| 13-KODE |
| PGF2a |
| 8,9-DiHETrE |
| 11,12-DiHETrE |

* Orange font, up in SO+f mice

| Dysregulated Metabolites in SO+f vs Viv chow in Conv and GF mice | | |
|---|---|---|
| 12-HETE * | 12-HEPE | 5, 6-DiHETrE |
| 2-LG | 14,15-DiHETE | 5-HEPE |
| 8,15-DiHETE | 14,15-DiHETrE | 9-HEPE |
| ALA | 15-HEPE | DHEA |
| LA | 17,18-DiHETE | EPA |
| 12,13-DiHODE | 19,20-DiHDoPA | EPEA |
| 12,13-DiHOME | 4-HDoHE | POEA |

| Dysregulated Metabolites in SO+f vs Viv chow in GF mice only |
|---|
| 14(15)-EpETrE |
| 6-keto PGF1a |
| AA |
| aLEA |
| DGLEA * |
| DHA |
| NA-Gly |
| PEA * |
| PGF3a * |

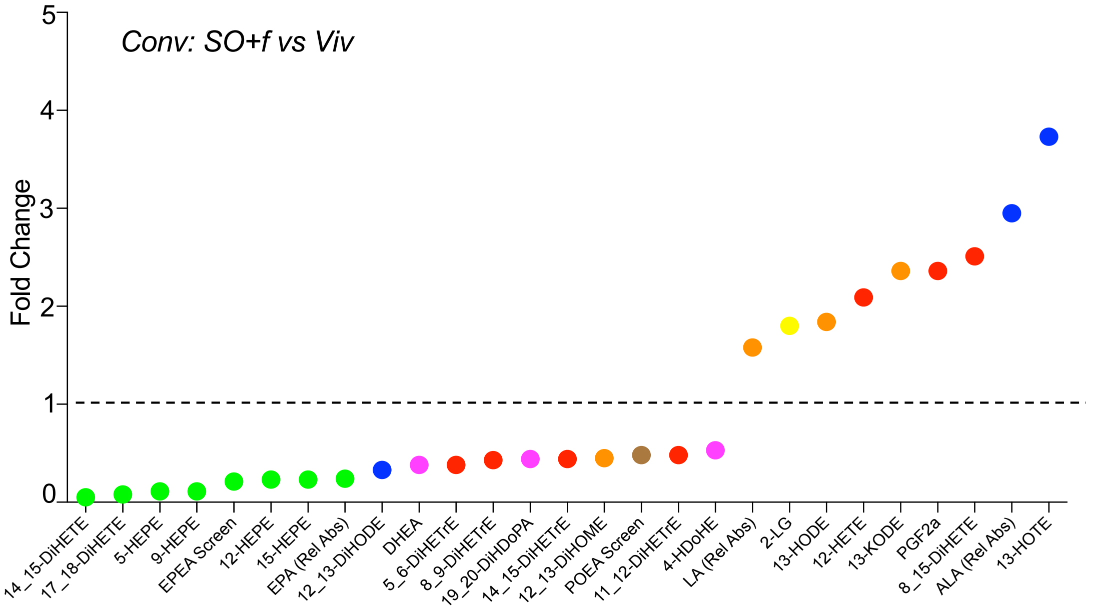


ALA, alpha linoleic acid
EPA, eicosapentenoic acid
LA, linoleic acid
AA, arachidonic acid
POA, palmitoleic acid
DHA, docosahexenoic acid
OA, oleic acid
LG, linoleoyl glycerol
PA, palmitic acid
DGLA, Dihomo-γ-linolenic acid

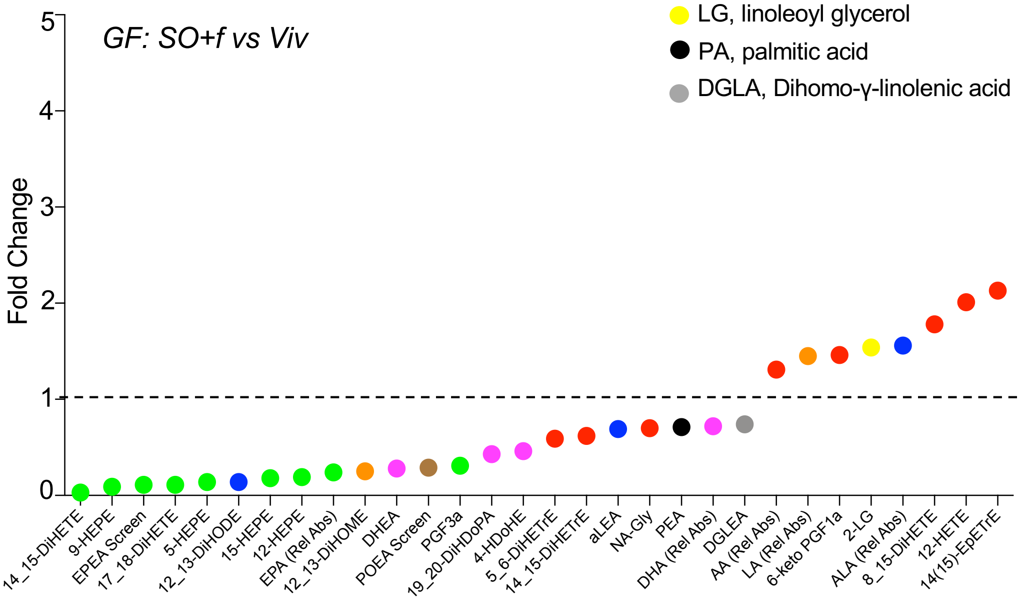


**Supplementary Figure 7. Comparison of effect of diet on oxylipin and endocannibinoid levels in IECs of conventional (Conv) and germ-free (GF) mice.** Table shows the significantly ($P \leq 0.05$) dysregulated metabolites in the different conditions.* also altered in SO-*m*AIEC grown with or without soybean oil in media: see Supplementary Table 4 and Supplementary Figure 5 for absolute levels. The graphs show the fold change in the indicated comparison with the metabolites color-coded by the parent compound.

| Dysregulated Metabolites in GF vs Conv in Viv chow only | | |
|---|---|---|
| 1-LG | 15, 16-DiHODE | 9-KODE* |
| 11-HETE* | 15(16)-EpODE | DHEA |
| 12,13-DiHODE | 2-AG | EPEA* |
| 12, 13-DiHOME | 2-LG | PEA* |
| 12(13)-Ep-9-KODE | 2-OG | PGF3a* |
| 12(13)-EpODE | 9, 10-DiHODE | POEA* |
| 13-HODE | 9-HODE | |
| 13-KODE | 9-HOTE | |

| Dysregulated Metabolites in GF vs Conv in Viv and SO+f | |
|---|---|
| AEA* | LA* |
| ALA* | LEA |
| aLEA | NA-Gly |
| DHA | OEA* |
| EPA* | PGF2a |
| F2 Isoprostanes | |

| Dysregulated Metabolites in GF vs Conv in SO+f only |
|---|
| 11, 12-DiHETrE |
| 17, 18-DiHETE |
| 5, 6-DiHETrE |
| 5-HEPE |
| 8, 9-DiHETrE |
| AA |
| NO-Gly* |

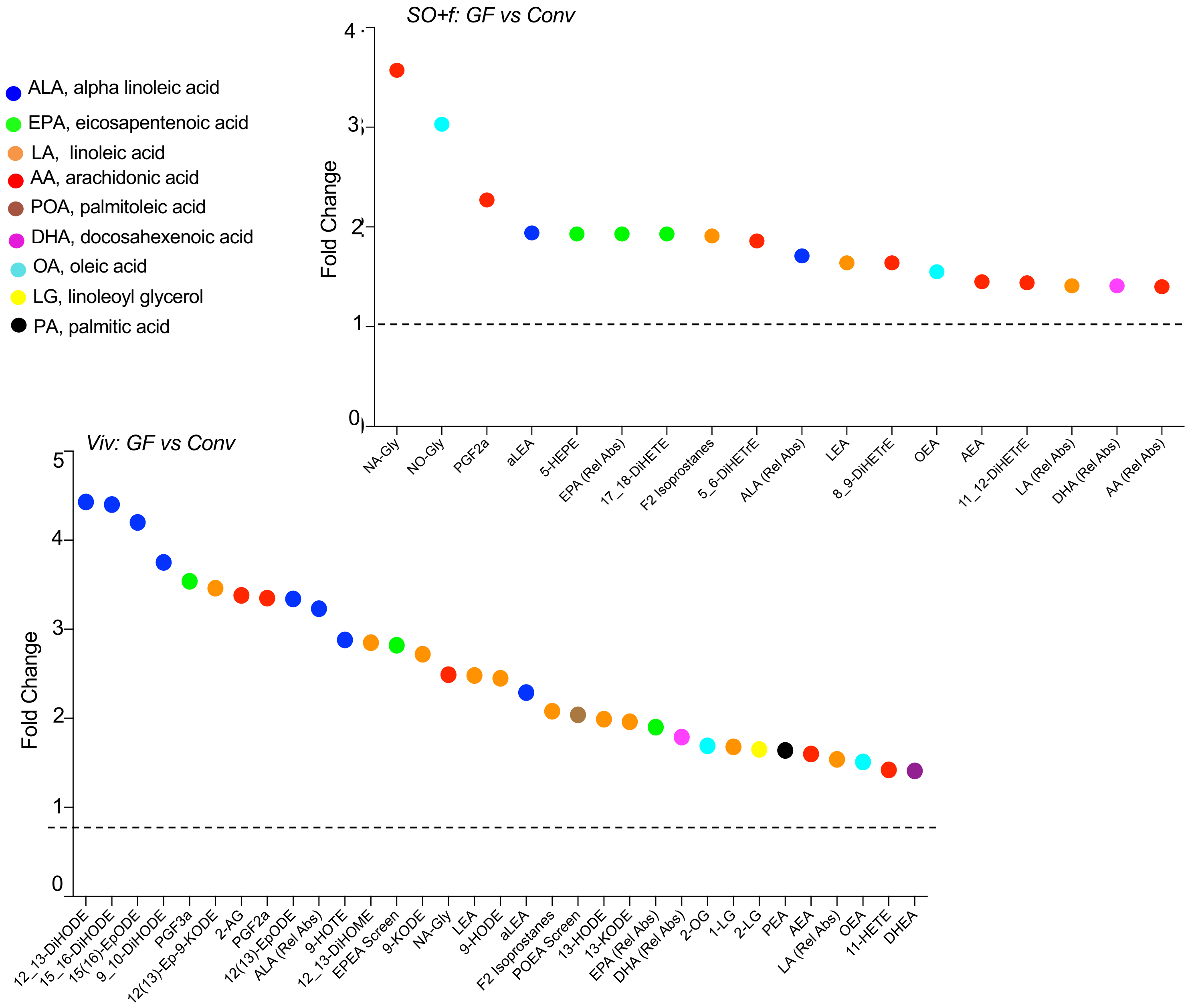


**Supplementary Figure 8. Comparison of effect of gut microbiota on oxylipin and endocannibinoid levels in IECs of conventional (Conv) versus germ-free (GF) mice.** Table shows the significantly (P≤ 0.05) dysregulated metabolites in the different conditions. * also altered in SO-*m*AIEC grown with or without soybean oil in media: see Supplementary Table 4 and Supplementary Figure 5 for absolute levels. The graphs show the fold change in indicated comparison with the metabolites color-coded by the parent compound.

*Distal colon: 12 weeks on diet, DSS treated*

WT Viv chow

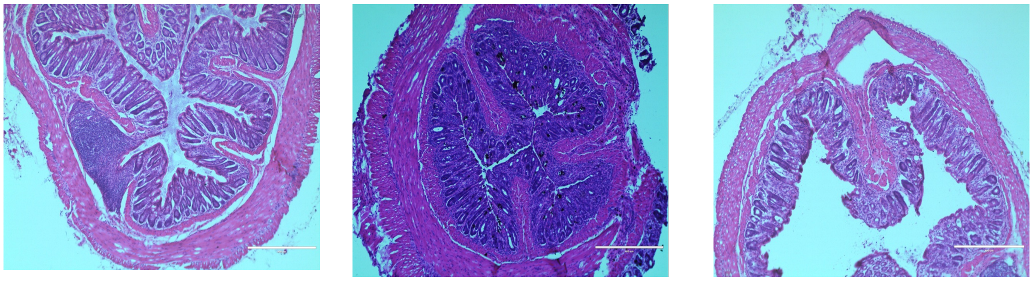

WT SO

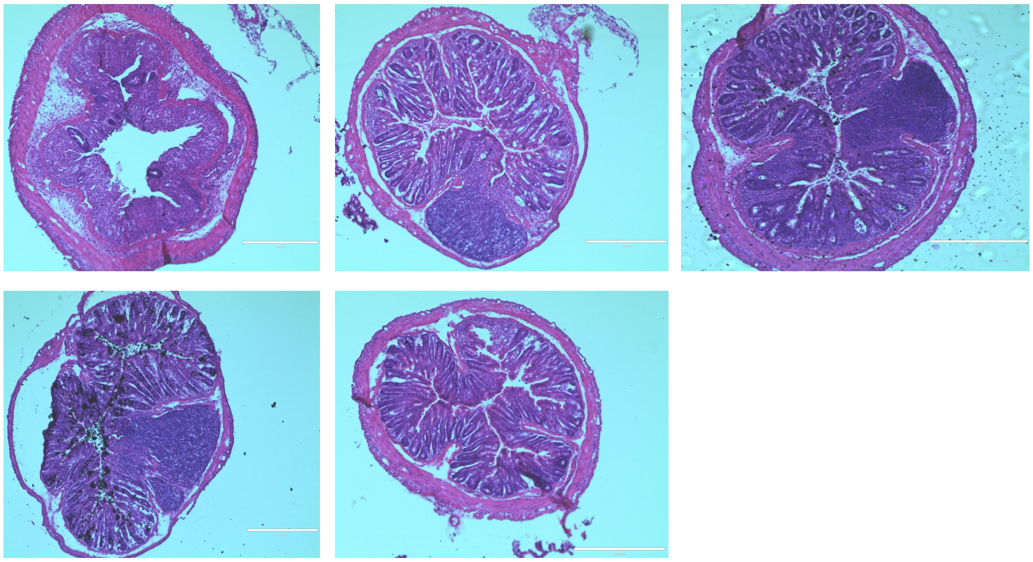

WT Plenish

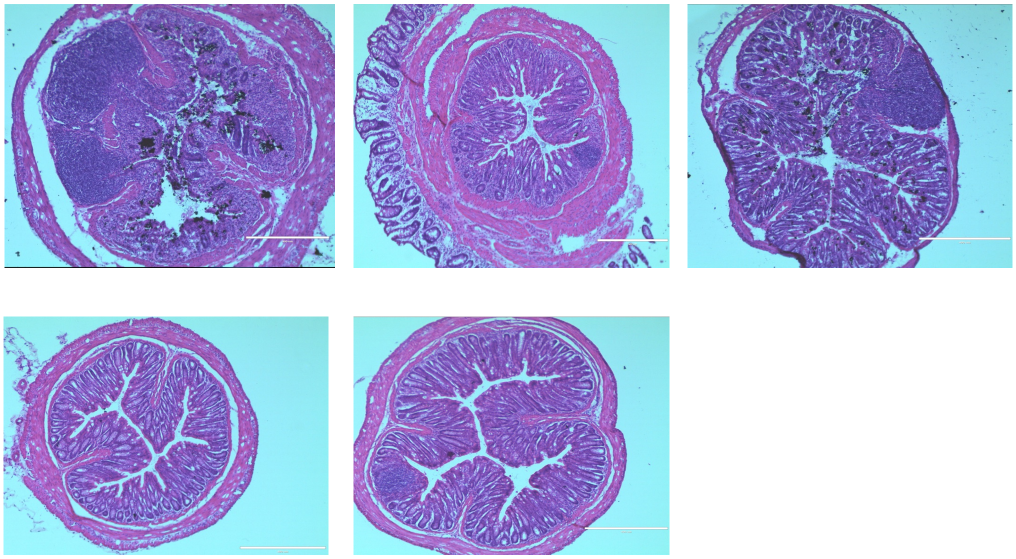

**Supplementary Figure 9. Distal colon histology in WT mice after DSS treatment.** H&E stains of distal colon from adult males on Viv chow, SO or low LA soybean oil (Plenish) diets for 12 weeks, treated with 2.5% DSS in drinking water for 6 days followed by 3 days recovery on water. N=4-6 per group. Calibration bar is 400μm.
