## Supplementary Table 1 for "Diet High in Soybean Oil Increases Susceptibility to Colitis in Mice"

**Supplementary Table 1.** Composition of diets and oils used in this study

| **Diet*:** | **Viv Chow** | **SO+f** | **SO** | **PL** | **OO** |
|---|---|---|---|---|---|
| Protein (gm%) | 23.9 | 16 | 19.5 | 19.5 | 19.5 |
| Carbohydrate (gm%) | 48.7 | 45.9 | 57.5 | 57.5 | 57.5 |
| Fat (gm%) | 5 | 14.9 | 18.2 | 18.2 | 18.2 |
| kcal/gm | 3.36 | 4.26 | 4.26 | 4.26 | 4.26 |
| Fat (kcal%) | 13.4 | 35 | 35 | 35 | 35 |
| LA (kcal%) | 3.3 | 18.6 | 19 | 2.6 | 4.5 |
| ALA (kcal%) | 0.27 | 3 | 3 | 0.67 | 0.2 |
| Oleic Acid (kcal%)** | 4.32 | 8 | 8 | 25.7 | 26.6 |
| Fiber (gm%)*** | 23.3 | 19.3 | 0 | 0 | 0 |
| **Source of Fat (gm%)** | **Viv Chow** | **SO+f** | **SO** | **PL** | **OO** |
| Porcine Animal Fat | 4.5 | 0 | 0 | 0 | 0 |
| Soybean Oil | 0 | 215 | 215 | 0 | 0 |
| Plenish Oil | 0 | 0 | 0 | 215 | 0 |
| Olive Oil | 0 | 0 | 0 | 0 | 215 |
| **Fatty acid composition of oils** | **Soybean** | **Plenish** | **Olive** | | |
| Lauric (12:0) | <0.05 | <0.05 | NA | | |
| Myristic (14:0) | 0.07 | <0.05 | NA | | |
| Palmitic (16:0) | 10.6 | 5.81 | 11.7 | | |
| Stearic (18:0) | 3.98 | 4.17 | 3.2 | | |
| Oleic (18:1) | 20.9 | 73.9 | 76.5 | | |
| Linoleic (18:2) | 52.9 | 7.42 | 6 | | |
| $\alpha$-Linolenic | 6.54 | 1.91 | 0.6 | | |

* Viv chow is Purina Test Diet 5001; all other diets formulated by Research Diets, Inc.
** Data only available for total monounsaturated fatty acids for Viv chow
*** Viv chow fiber composed of cellulose, hemi-cellulose and lignin; SO+f fiber comprised of cellulose + inulin
