## Supplementary Table 2 for "Diet High in Soybean Oil Increases Susceptibility to Colitis in Mice"

**Supplementary Table 2.** Scoring criteria for Disease Activity Index (DAI).

| Score* | Weight Loss (%) | Colon Length/ Weight Ratio | Hemoccult | Morphological changes |
|---|---|---|---|---|
| 0 | 0 to 5 | <40 | negative | normal |
| 1 | 5 to 10 | 40 to 45 | very faint | inflamed |
| 2 | 10 to 15 | 45 to 50 | faint | polyps/nodules |
| 3 | 15 to 20 | >50 | moderate | Prolapsed rectum |
| 4 | >20 | | excessive | > one change |

* The DAI per mouse was calculated by totaling the scores for each of the four criteria.
